# Fine-Grained and Reliable Cell Classification from Pathology Images

**DOI:** 10.64898/2026.10.20.760347

**Authors:** Sevda Öğüt, Arpit Merchant, Amirhossein Vahidi, Dabin Jeong, Sebastian Birk, Adam Boxall, Adib Miraki Feriz, Marta Rosa Sallese, Yinshui Chang, Pascal Frossard, Dorina Thanou, Mo Lotfollahi

**Author notes:** Work done while visiting the Wellcome Sanger Institute. Correspondence. Co-last authorship (Dorina Thanou, Mo Lotfollahi).

## Abstract

Molecular assays such as spatial transcriptomics facilitate fine-grained identification of cell types for analyzing tumor microenvironments. However, they are expensive to acquire in routine clinical practice compared to hematoxylin and eosin (H&E) images. With molecular guidance, such images can distinguish cell lineages, yet often fail to resolve closely related subtypes. To overcome this, we introduce **CellBridge**, a multi-modal contrastive learning framework pretrained on 16M cells with paired H&E and spatial profiles, for transferring molecular information into morphological representations. CellBridge further incorporates a Cell-Ontology-aware conformal prediction module that assigns labels at different levels of specificity: leaf-level cell types when morphology alone provides sufficient evidence, and broader ancestor categories when distinguishing among more specific cell types would require molecular information. Across in-domain and out-of-domain benchmarks, CellBridge outperforms leading pathology foundation models in macro F1 by up to 16 percentage points, with the largest gains on the most fine-grained label spaces. It consistently achieves smaller prediction sets, while reducing class-wise coverage gaps. We envision CellBridge as a step toward extracting cellular information that currently requires costly molecular assays from routinely acquired pathology images, while explicitly identifying where such assays remain necessary. Our code is available at https://github.com/Lotfollahilab/cellbridge.

## 1 Introduction

Cells are core components of the tumor microenvironment (TME) which governs diagnosis, prognosis, and treatment in cancer (Wang et al., 2023; Peyraud et al., 2025). Hematoxylin and eosin (H&E) staining is the most widely available readout of TMEs, acquired for nearly every patient and digitized at scale. Yet, identifying fine-grained cell types from H&E remains difficult for two reasons. First, subtypes within a lineage, such as T and B lymphocytes, share highly similar morphology. Second, cells from distinct lineages can appear alike, as with macrophages infiltrating tumor nests.

Spatial transcriptomics (ST) provides a complementary view that resolves many of these ambiguities. Imaging-based platforms such as Xenium measure gene expression in individually segmented cells, and molecular profiles separate cell types that morphology alone cannot. However, the cost and throughput of ST preclude routine clinical use. Emerging public datasets that pair H&E with single-cell ST, such as HEST-1k (Jaume et al., 2024), offer a middle path: a model can learn from the molecular modality during training and then operate on H&E alone at inference. Most existing multi-modal approaches, however, align images with spot-level expression that aggregates signals across multiple cells (Hemker et al., 2026; Xiang et al., 2026; Huang et al., 2025), which obscures precisely the cell-level differences needed for fine-grained phenotyping.

Molecular supervision can reduce morphological ambiguity, but it cannot remove it entirely, since some distinctions are simply not visible in an H&E image. A central question is therefore not only which cell type a model predicts, but how far along the cell-type hierarchy H&E can reliably predict. Current pathology models, both unimodal and multi-modal, cannot answer this, because they treat cell types as independent, flat labels and ignore the taxonomic relationships between them, which are available in the Cell Ontology (Tan et al., 2026). In transcriptomics, leveraging these relationships has been shown to yield more biologically meaningful cell representations (Yuan et al., 2024). For H&E, the ontology additionally offers a natural scale on which to express the limit of predictability. For instance, a model may reliably identify a cell as a lymphocyte, yet be unable to determine from morphology whether it is a T cell or a B cell. Reporting *lymphocyte* in this case is both correct and informative. It states what the image supports and signals that resolving the cell further requires additional data, such as ST or multiplex imaging. A flat classifier instead commits to a single finegrained label and is confidently wrong in precisely these cases.

To address these limitations, we propose CellBridge, a framework for fine-grained and reliable phenotyping of single cells from H&E images. CellBridge operates at two levels. (1) At the cross-modal level, it transfers molecular identity into morphological representations by aligning cell-centered H&E crops from a pathology foundation model with embeddings from a frozen single-cell ST foundation model, using a contrastive objective and parameter-efficient adaptation on 16M paired Xenium cells (Section 2.1). This sharpens distinctions that morphology alone cannot make, while requiring only images at inference. (2) At the ontology level, a Cell-Ontology-Fallback (COF) module converts point predictions into biologically coherent prediction sets via conformal prediction calibrated within Cell Ontology branches (Section 2.2). For each cell, CellBridge reports the most specific ontology term supported by the image, with coverage guaranteed within each lineage. This makes the predictive limit of H&E explicit by returning leaf-level phenotypes where morphology suffices, and broader ancestors where molecular measurements are needed to resolve the cell.

We evaluate CellBridge against six patch-level, cell-level, and multi-modal foundation models on 14 held-out HEST sections and four external datasets. CellBridge improves macro F1 by up to 16 percentage points over the strongest baseline, with gains increasing as label spaces become more fine-grained (Section 3.2). Under COF, it ranks best or second-best in prediction-set size and singleton rate on every dataset (Section 3.3), and branch-conditional calibration reduces class-wise coverage gaps up to fivefold relative to standard conformal prediction (Section 3.4). Finally, expert biological review shows that low reliability scores localize to biologically ambiguous cells, such as macrophages infiltrating tumor clusters and cells at the myoepithelial–tumor interface (Section 3.5).

Our contributions are as follows:

- We introduce CellBridge, which aligns morphology and gene expression at single-cell resolution through multi-modal contrastive pretraining on 16M paired cells, enabling fine-grained phenotyping from H&E alone.
- We propose a Cell-Ontology-Fallback module, which uses the Cell Ontology to determine, for each cell, the finest phenotype that can be reliably predicted from H&E, with per-lineage coverage guarantees, thereby indicating where additional molecular data is required.
- We show that CellBridge improves both the accuracy of fine-grained classification and the reliability of ontology-aware categorization across in-domain and out-of-domain benchmarks, with low-confidence predictions localizing to biologically ambiguous cells.

## 2 Cell Bridge

CellBridge operates at two levels: cross-modal representation learning and ontology-aware prediction. We first describe how molecular supervision guides learning single-cell image representations, then explain our COF module, which uses the Cell Ontology to adapt prediction granularity.

### 2.1 Molecularly guided single-cell image representations

We aim to identify cellular image features that are predictive of a cell’s molecular identity. To this end, we leverage paired H&E images and single-cell ST measurements to supervise the image encoder toward representations that are predictive of cellular transcriptional state. Formally, let 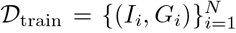 denote our training dataset, where *N* is the total number of cells, *I*_*i*_ is an H&E crop centered on cell *i*, and *G*_*i*_ is its corresponding gene expression profile.

As illustrated in Figure 1, our cross-modal alignment leverages two pretrained foundation models. A pathology foundation model *f*_PFM_ encodes the image crop *I*_*i*_, while a single-cell foundation model *f*_scFM_ encodes the gene expression *G*_*i*_. To bridge these modalities efficiently, we freeze *f*_scFM_ and the base parameters of *f*_PFM_, adapting only its final layers via Low-Rank Adaptation (LoRA) (Hu et al., 2022). Specifically, each adapted linear layer within the image encoder is defined as

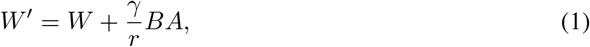

where *W* is the frozen weight matrix, *A* and *B* are trainable low-rank matrices, *r* is the rank of the update, and *γ* is a scaling factor. A trainable projection head *P* then maps the image features to the dimensionality of the transcriptomic embeddings. The normalized representations for alignment are

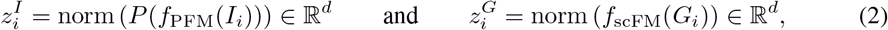

where norm(*v*) = *v/* ∥*v*∥_2_. For a batch of *M* paired cells within a tissue section, we optimize a symmetric InfoNCE objective (Radford et al., 2021), treating matched image–expression pairs as positives and other pairs within the batch as negatives. Hence, the alignment loss becomes

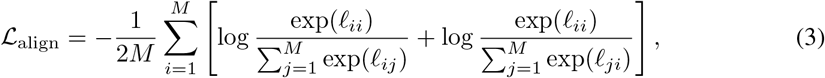

with 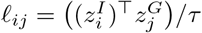, where *τ* is the temperature. During this multi-modal contrastive pretraining, only the LoRA parameters and projection head are optimized, encouraging the image representation to capture molecular information associated with the target cell.

**Figure 1.**
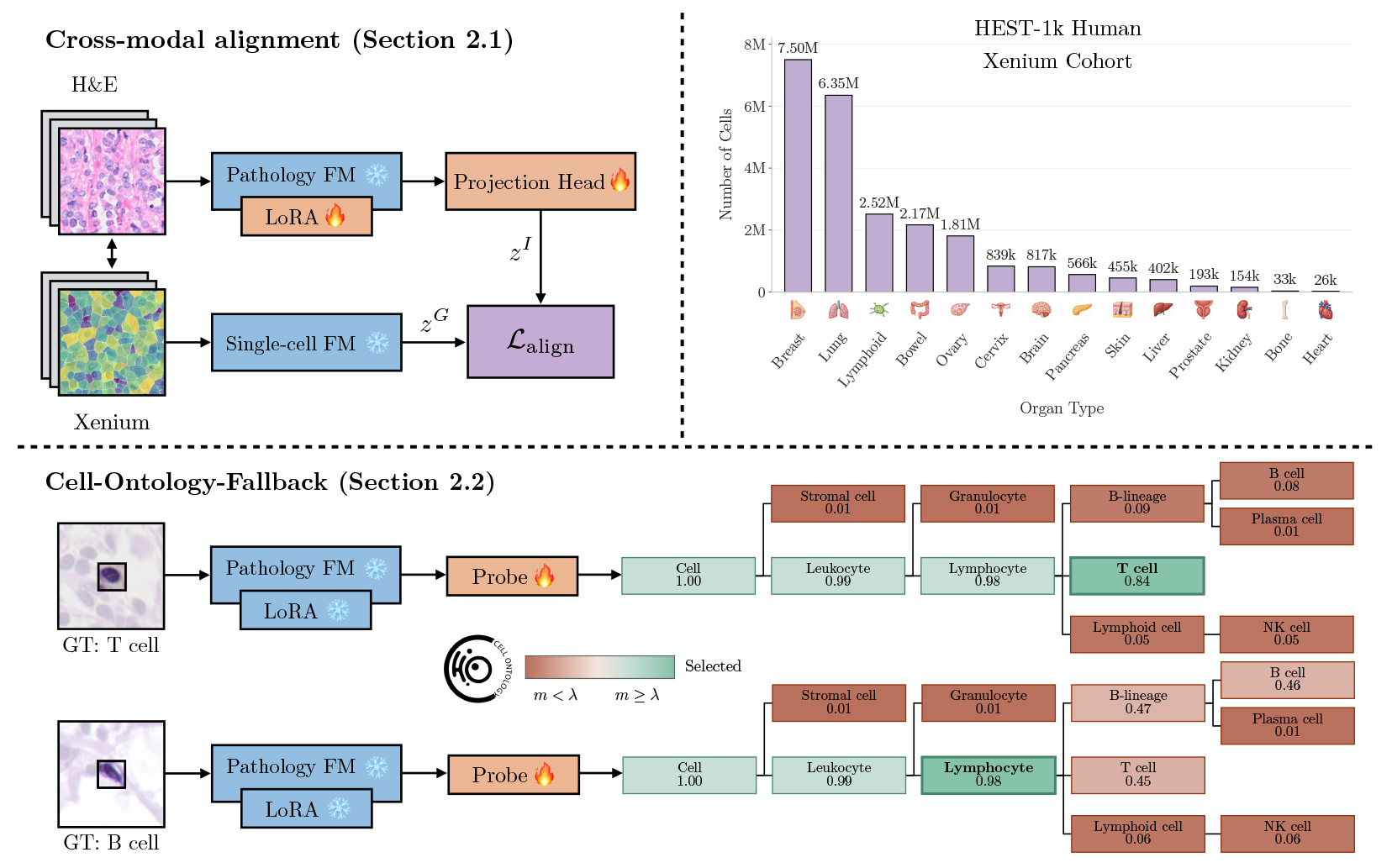
Overview of our framework. *Top:* During pretraining, CellBridge aligns representations from paired H&E images and gene expression transcripts from the Xenium HEST-1k human cohort. It adapts a pathology foundation model via LoRA and a trainable projection head to learn the embedding space of a single-cell spatial foundation model using a multi-modal contrastive objective. *Bottom:* At evaluation, CellBridge operates solely on images. Its Cell-Ontology-Fallback module dynamically adjusts prediction granularity based on a *λ*. Predictions with probability mass *m* above *λ* yield fine-grained, leaf-level phenotypes, whereas those below fall back to coarser common ancestors, ensuring reliability guarantees. Values beneath ontology terms indicate their assigned probability mass. Snowflake and fire icons denote frozen and trainable components, respectively.

### 2.2 Conformal prediction with Cell-Ontology-Fallback

After pretraining, we discard both *f*_scFM_ and *P*, retaining only the adapted image encoder, denoted by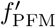. We then extract features directly from image crops centered on precomputed cell coordinates^1^ and fit a lightweight probe *f*_θ_ using labeled training samples.

While *f*_θ_ predicts fine-grained cell types, not every cell can be reliably resolved at the same level of specificity from H&E. We therefore introduce *Cell-Ontology-Fallback* (COF), which combines conformal prediction (CP) with the Cell Ontology to determine the most specific phenotype supported by the model for each cell. This module converts a classifier into a set-valued predictor using heldout labeled data (Vovk et al., 2005). By calibrating a nonconformity threshold on this held-out set, the model dynamically returns either a specific phenotype or a coarser set of plausible candidates.

#### Calibration setup and coverage target

We reserve a subset 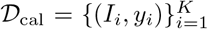 for calibration, keeping the image encoder and lightweight probe fixed. For a test crop *I*_test_, our goal is to return a biologically coherent set *C* (*I*_test_) ⊆*Y* that respects the hierarchical lineages and satisfies

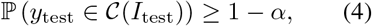

for a user-defined miscoverage level *α* ∈[0, 1] and under the exchangeability of calibration and test cells. To achieve this, our COF module defines (i) a Cell-Ontology-based nonconformity score (Section 2.2.1), and (ii) a calibration rule within biological branches (Section 2.2.2), detailed below.

##### Algorithm 1

Cell-Ontology-Fallback module

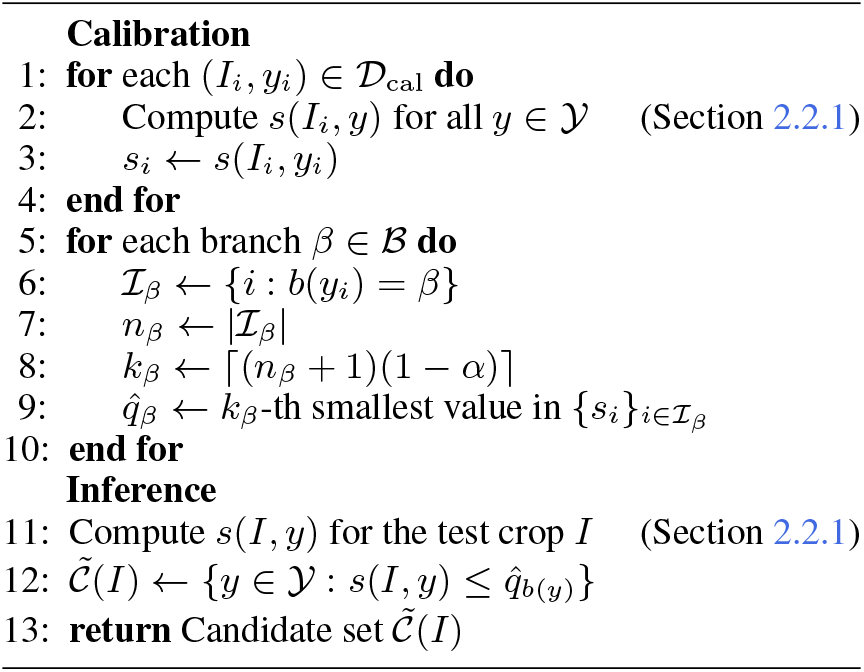

#### 2.2.1 Biologically-informed prediction sets

A nonconformity score *s*(*I*_i_, *y*) measures the disagreement between an image crop centered on cell *i* and a candidate phenotype *y*, with lower values indicating greater agreement. Following Corbetta et al. (2026), we define this score through a nested family of prediction sets informed by the Cell Ontology (Tan et al., 2026), incorporating biological relationships into the possible outputs.

In particular, we construct a task-specific directed acyclic graph (DAG) by retaining the ontology terms corresponding to the evaluated phenotypes and their ancestors. For a cell *i*, we start from the highest-probability phenotype 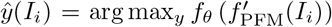 and consider its leaf node and ancestors as candidate output categories. Let *L*(*v*) ⊆*Y* denote the phenotype leaves descending from node *v*. Each candidate node receives the aggregate probability mass of its descendant phenotypes

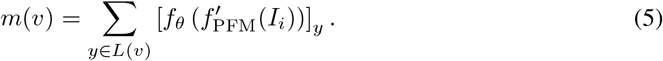

For a threshold *λ* ∈ [0, 1], we select the candidate node with the smallest mass satisfying *m*(*v*) ≥ *λ*, resolving ties deterministically. We then construct the set *S*_*λ*_(*I*_i_) to be the descendant phenotypes of this selected node, together with those of all considered nodes whose mass is below *λ*. This union ensures that the sets remain nested as *λ* increases, including when the ontology contains multiple inheritance. Lastly, we define *s*(*I*_i_,*y*) as the smallest evaluated threshold *λ* at which phenotype *y* enters *S*_*λ*_(*I*_i_). Thus, lower scores identify phenotypes included at smaller thresholds, whereas higher scores require further expansion of the prediction set along the ontology.

#### 2.2.2 Branch-conditional conformal calibration

Given these scores, calibration determines the thresholds used to admit candidate phenotypes into the prediction set. Global calibration uses a single threshold and guarantees marginal coverage, but can conceal undercoverage of individual phenotypes. Class-conditional calibration addresses this limitation, yet scarce calibration examples for rare subtypes can lead to overly conservative thresholds and larger prediction sets. Motivated by this trade-off and subgroup-conditional Mondrian CP (Vovk et al., 2003), we calibrate scores separately within predefined ontology branches.

A fixed mapping *b* : *Y* → *B* assigns each phenotype to a unique configured ancestor branch, ensuring disjoint calibration groups. For each branch *β* ∈ *B*, let *I*_*β*_ = {*i* : *b*(*y*_i_) = *β* } collect its *n*_β_ calibration cells. We evaluate each cell’s nonconformity score against its true phenotype, *s*_i_ = *s*(*I*_i_, *y*_*i*_), and compute the branch-specific threshold

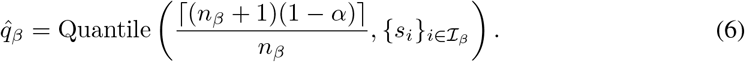

At inference, each phenotype is admitted according to the calibrated threshold of its own branch:

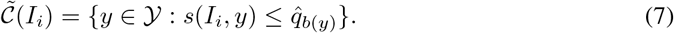

Under exchangeability within each branch, this yields, for every *β B*

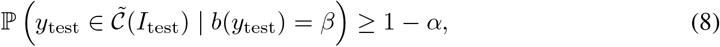

which is a guarantee that holds within each biological group and consequently implies Equation 4. However, the branch-calibrated candidate set need not coincide with the descendant labels of a single ontology node, allowing it to capture ambiguity that does not align with the ontology hierarchy. We therefore apply a closure operation. Specifically, for a nonempty candidate set, we select the node *v*^⋆^(*I*_*i*_) that satisfies

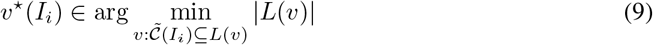

break ties deterministically, and return *C*(*I*_i_) = *L*(*v*^⋆^ (*I*_i_)). Since this operation only enlarges the candidate set, it preserves the branch-conditional coverage guarantee. Finally, we summarize prediction specificity through a *reliability score* for each cell *i*

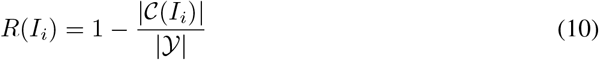

where smaller prediction sets correspond to higher scores. In practice, model outputs with low scores can be mapped to the lowest common ancestor of their constituent labels in the Cell Ontology, yielding a broader category, as conveyed in Figure 1.

## 3 Experiments

Through extensive empirical evaluation, we answer the following research questions: *RQ1:* Does multi-modal pretraining improve fine-grained cell phenotyping from H&E on unseen sections? *RQ2:* How well do the learned representations generalize across datasets and phenotype taxonomies? *RQ3:* Can the COF module balance phenotypic specificity with predictive reliability?

### 3.1 Setup

#### Datasets and splits

We utilize the human Xenium cohort of HEST-1k (23.8M cells), which pairs H&E images with singlecell spatial transcriptomic profiles, for pretraining and cross-section evaluation (Jaume et al., 2024). Pretraining and model selection use 59 and 6 sections, respectively, whereas our test set consists of 14 annotated sections spanning breast, ovary, and pancreas, held out from both stages. Within each unseen sample, we follow Hizmi et al. (2026) and reserve the bottom 20% of cells for testing. For downstream analysis, we use the remaining 80% for three-fold cross-validation. For conformal prediction, we further partition these cells into training and calibration sets comprising 64% and 16% of the total cells, respectively, as shown in Appendix Figure 5. In addition, we assess external generalization on four datasets with complementary annotation sources and both fine-grained and coarse phenotype taxonomies. Namely, breast cancer nuclei from NuCLS (Amgad et al., 2022), the multi-organ PanNuke dataset (Gamper et al., 2020), colon cancer images from Lizard (Graham et al., 2021), and CODEX-derived colon carcinoma from PhenoCell, where we evaluate both the full taxonomy and the immune subset. We use fixed training, validation, and test partitions, following PathoCellBench (Lüscher et al., 2025). Dataset statistics are summarized in Table 1 with additional details in Appendix A.1.

**Table 1:**
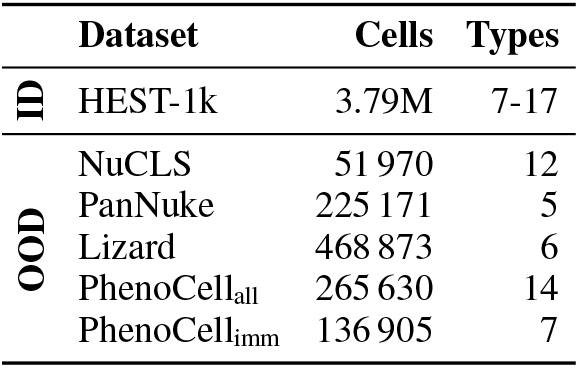
Evaluation datasets.

#### Pretraining of CellBridge

We instantiate the pathology encoder with UNI2-h (Chen et al., 2024) and the single-cell encoder with TERRA-112M (Birk et al., 2026), which produce 1536- and 384-dimensional embeddings, respectively. We insert LoRA adapters into the fused query, key, value, and output projections of the final four attention blocks of UNI2-h, using rank *r* = 8 and scaling factor *γ* = 16 in Equation 1. Moreover, we set the contrastive loss temperature to *τ* = 0.07 and train for one epoch using AdamW (Loshchilov & Hutter, 2019) with batches of 16 paired image– expression observations. The best checkpoint is selected with respect to the lowest validation loss. Implementation details along with time and memory requirements are provided in Appendix A.3.

#### Baseline models

We compare CellBridge with six foundation models spanning three categories. *Patch-level FMs* learn representations from pathology tiles and include UNI2-h (Chen et al., 2024) and Virchow2 (Zimmermann et al., 2024), selected for their strong performance across multiple benchmarks (Tchokponhoue et al., 2026; Campanella et al., 2025). *Cell-level FMs* learn representations from single-cell images and consist of VOLTA (Nakhli et al., 2024) and LEMON (Chadoutaud et al., 2026). *Multi-modal FMs* incorporate molecular supervision through paired data and involve PAST (Yang et al., 2025) and SpatialWhisperer (Schaefer et al., 2026). All encoders remain frozen during downstream evaluation, and following Jaume et al. (2024), we reduce the patch-level FM and CellBridge embeddings to 256 dimensions using PCA fitted only on training data. For the remaining models, we retain the recommended native dimensionality of embeddings. Further details are documented in Appendix A.2.

#### Evaluation metrics

For downstream analysis, we fit both a linear probe and a cosine 20-nearest-neighbor probe on identical splits. We assess performance with macro F1, balanced accuracy, and macro AUROC on our ID and OOD datasets. For the former, we take the mean and standard deviation across held-out HEST sections, whereas for the latter, we report the mean and standard error across 1000 grouped-bootstrap replicates of the fixed test sets. Furthermore, for conformal prediction, we similarly follow the same linear probe strategy, while targeting *α* = 0.1 miscoverage in Equation 4 as in Ding et al. (2023). Our metrics for this part include the mean set size, namely the average number of phenotype labels in the returned sets, and the singleton rate, meaning the proportion of test cells receiving a set containing exactly one phenotype. Additional details are provided in Appendix A.3.

### 3.2 Representation transfer to unseen sections and external datasets

#### CellBridge representations transfer to unseen sections

Figure 2 compares macro F1 on 14 held-out HEST sections under the spatial split shown in Appendix Figure 5. Addressing *RQ1*, CellBridge achieves the highest performance with both probes, reaching approximately 50% macro F1 on sections excluded from pretraining, despite substantial class imbalance and label spaces ranging from 7 to 17 phenotypes as seen in Table 1. PAST is the strongest baseline, consistent with the potential benefits of pretraining on paired imaging and molecular data.

**Figure 2.**
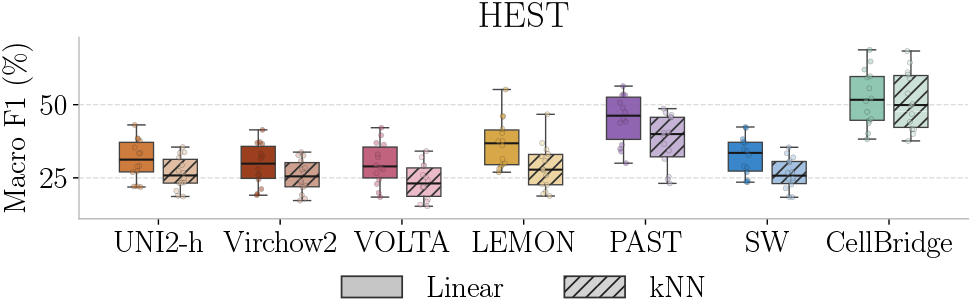
Cross-section transferability of CellBridge. Macro F1 (%) across 14 held-out HEST sections for linear (solid) and kNN (hatched) probes. Boxes show medians and interquartile ranges, and dots denote individual sections.

Moreover, CellBridge exhibits a notably smaller gap between linear and kNN probing than the baselines. In fact, its kNN probe surpasses the linear probes of most competing methods. This suggests that molecularly guided adaptation better reflects cell identity, allowing a simple similarity-based classifier to approach the performance of a trained linear probe. Together, these results support the transferability of CellBridge representations to unseen tissue sections. The corresponding balanced accuracy and macro AUROC plots show similar trends and are provided in Appendix A.4.

#### CellBridge embeddings generalize across external datasets

Table 2 demonstrates that the molecularly informed representations learned by CellBridge generalize to five external H&E-only tasks, addressing *RQ2*. Our method achieves the highest macro F1 in nine of ten dataset-probe configurations, outperforming patch-level, cell-level, and multi-modal FMs. Under linear probing, the largest gains over the second-best method occur on NuCLS (+16.0 pp) and PhenoCell_all_ (+12.0 pp). Notably, these cohorts feature the most complex and granular cell types, underscoring the model’s capacity to resolve challenging label spaces, as in Table 1 and Appendix Table 7.

**Table 2:** OOD generalization across external datasets. Macro F1 (%) scores comparing Cell-Bridge against baseline FMs using both linear and kNN probes. Values denote the mean and standard error computed across 1000 grouped-bootstrap replicates of the fixed test sets. SW stands for SpatialWhisperer. Bold and underline indicate the best and second-best per row, respectively.

|  |  | Patch-level FMs |  | Cell-level FMs |  | Multi-modal FMs |  | Ours |
| --- | --- | --- | --- | --- | --- | --- | --- | --- |
|  |  | UNI2-h | Virchow2 | VOLTA | LEMON | PAST | SW | CellBridge |
| Linear Probe | NuCLS | 23.7 ± 1.37 | 25.1 ± 2.83 | 22.7 ± 2.16 | 24.0 ± 2.35 | 23.7 ± 2.12 | 24.2 ± 2.16 | **41.1 ± 2.82** |
| | PanNuke | 72.5 ± 1.41 | 72.1 ± 1.52 | 56.9 ± 1.00 | 67.7 ± 1.10 | 74.3 ± 1.18 | 73.1 ± 1.20 | **79.1 ± 1.11** |
| | Lizard | 35.7 ± 1.28 | 35.6 ± 1.81 | 59.3 ± 1.38 | **69.6 ± 0.92** | 47.1 ± 1.34 | 37.0 ± 0.85 | <u>65.4 ± 1.36</u> |
| | PhenoCell<sub>all</sub> | <u>25.1 ± 2.07</u> | 24.7 ± 2.07 | 15.4 ± 0.97 | 22.0 ± 1.34 | 23.1 ± 1.83 | 24.2 ± 1.91 | **37.1 ± 3.24** |
| | PhenoCell<sub>imm</sub> | <u>37.3 ± 4.39</u> | 33.9 ± 4.03 | 24.1 ± 1.37 | 31.6 ± 1.93 | 29.5 ± 2.14 | 32.0 ± 3.11 | **41.2 ± 3.31** |
| kNN | NuCLS | 18.5 ± 0.97 | 21.0 ± 1.78 | 21.2 ± 1.09 | <u>23.7 ± 1.60</u> | 18.0 ± 1.47 | 21.2 ± 1.12 | **31.7 ± 2.02** |
| | PanNuke | 72.9 ± 1.21 | 73.3 ± 1.17 | 47.3 ± 0.61 | 65.8 ± 1.20 | 75.8 ± 1.11 | 73.0 ± 1.24 | **80.4 ± 0.99** |
| | Lizard | 28.7 ± 1.08 | 28.0 ± 1.02 | 41.6 ± 0.86 | <u>50.1 ± 1.43</u> | 33.6 ± 0.97 | 29.7 ± 1.10 | **60.9 ± 1.52** |
| | PhenoCell<sub>all</sub> | <u>22.4 ± 1.80</u> | 19.9 ± 1.41 | 13.3 ± 0.85 | 16.8 ± 1.21 | 18.9 ± 1.18 | 21.7 ± 1.73 | **35.0 ± 2.75** |
| | PhenoCell<sub>imm</sub> | <u>30.2 ± 3.50</u> | 27.1 ± 2.47 | 21.3 ± 1.60 | 25.4 ± 1.84 | 24.4 ± 1.99 | <u>30.3 ± 3.20</u> | **38.0 ± 2.99** |

#### CellBridge maintains its advantage across label granularities

We evaluate CellBridge against the baselines using the NuCLS dataset, as it is the only dataset in our study that provides explicitly nested label sets. These sets span coarse, medium, and detailed groupings with 5, 8, and 12 classes, respectively. As shown in Figure 3, macro F1 decreases for all methods as the label space becomes more detailed. Nevertheless, CellBridge consistently achieves the highest performance, with its advantage over the strongest baseline increasing from approximately +10.0 pp at the coarse level to +16.0 pp at the most detailed level.

**Figure 3.**
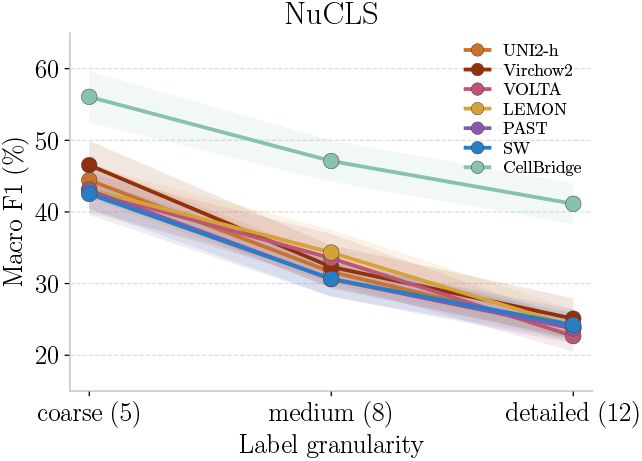
Strong performance of Cell-Bridge across varying label spaces. Reporting conventions follow Table 2.

### 3.3 Biologically-informed conformal prediction sets

#### CellBridge improves prediction-set efficiency

Table 3 presents the results comparing CellBridge against the six baselines under the same COF module at *α* = 0.1, so that differences in set size and singleton rate reflect the informativeness of the underlying representations. We observe the following that address *RQ3*. First, CellBridge ranks best or second-best on every dataset for both metrics. This clearly shows its consistency, especially given that no baseline attains the same ranking across datasets. In particular, PAST is competitive on HEST and PanNuke but returns the largest sets and only 11.5% singletons on NuCLS. Similarly, LEMON is strong on Lizard but markedly weaker on PanNuke. Second, gains are most pronounced on PanNuke and Lizard, where CellBridge reduces the mean set size by 0.37 and 0.64 labels and raises the singleton rate by 14.8 and 11.3 points over the best competitor, respectively. These gains verify that CellBridge also improves prediction specificity in coarser label spaces seen in Table 1 and Appendix Table 8, extending its benefits beyond fine-grained labels. In addition, as seen in Table 2, LEMON obtains the highest linear probing macro F1 on Lizard, yet requires larger sets to reach the same coverage, indicating conformal efficiency is not implied by point-prediction accuracy. Finally, both PhenoCell variants remain difficult for all models, with sets spanning roughly two-thirds of the label space and singleton rates below 25%. This confirms that our COF module returns coarser ontology terms, rather than unsupported leaf-level predictions, which is the intended behavior.

**Table 3:** Conformal prediction performance. Mean set size (↓) and singleton rate (↑) at a target coverage of 1 − *α* = 0.9 under our Cell-Ontology-Fallback module. Values report the mean and standard error, with the latter derived from the standard deviation across the ID sections and estimated via 1000 grouped-bootstrap replicates for OOD datasets. SW denotes SpatialWhisperer. Bold and underline indicate the best and second-best per row, respectively. Corresponding empirical coverage is reported in Appendix Table 12, which shows all methods successfully attain the target.

| Dataset |  | Patch-level FMs |  | Cell-level FMs |  | Multi-modal FMs |  | Ours |
| --- | --- | --- | --- | --- | --- | --- | --- | --- |
|  |  | UNI2-h | Virchow2 | VOLTA | LEMON | PAST | SW | CellBridge |
| Mean Set Size (↓) | | | | | | | | |
| ID | HEST | 7.19 ± 0.49 | 7.46 ± 0.57 | 7.52 ± 0.64 | 7.05 ± 0.57 | <u>5.85 ± 0.39</u> | 7.19 ± 0.55 | **5.05 ± 0.43** |
| OOD | NuCLS | 3.03 ± 0.10 | <u>2.78 ± 0.12</u> | 3.44 ± 0.15 | 3.22 ± 0.13 | 4.02 ± 0.08 | 3.12 ± 0.09 | **2.72 ± 0.12** |
| | PanNuke | 2.07 ± 0.02 | <u>2.06 ± 0.02</u> | 2.74 ± 0.01 | 2.26 ± 0.01 | <u>1.93 ± 0.02</u> | 2.04 ± 0.02 | **1.56 ± 0.01** |
| | Lizard | 4.15 ± 0.18 | 4.79 ± 0.20 | 3.67 ± 0.09 | <u>3.09 ± 0.09</u> | 3.37 ± 0.13 | 4.50 ± 0.15 | **2.45 ± 0.10** |
| | PhenoCell<sub>all</sub> | <u>8.29 ± 0.51</u> | 8.38 ± 0.57 | 9.37 ± 0.35 | 8.84 ± 0.31 | 8.47 ± 0.42 | 8.45 ± 0.49 | **7.70 ± 0.40** |
| | PhenoCell<sub>imm</sub> | 4.74 ± 0.37 | 4.74 ± 0.34 | 5.29 ± 0.22 | 5.27 ± 0.11 | **4.61 ± 0.27** | 4.68 ± 0.32 | <u>4.64 ± 0.28</u> |
| Singleton Rate (↑) | | | | | | | | |
| ID | HEST | 18.1 ± 2.84 | 15.6 ± 3.10 | 16.7 ± 3.14 | 20.4 ± 2.92 | <u>31.6 ± 3.10</u> | 18.2 ± 2.81 | **40.5 ± 3.68** |
| OOD | NuCLS | 33.4 ± 3.38 | **42.6 ± 3.37** | 30.7 ± 3.30 | 35.2 ± 4.12 | 11.5 ± 2.30 | 29.4 ± 3.12 | <u>39.5 ± 4.46</u> |
| | PanNuke | 53.7 ± 0.99 | 54.3 ± 0.96 | 33.4 ± 0.45 | 50.1 ± 0.55 | <u>60.6 ± 0.66</u> | 54.9 ± 0.86 | **75.4 ± 0.51** |
| | Lizard | 35.8 ± 3.91 | 22.9 ± 4.06 | 43.2 ± 1.95 | <u>54.5 ± 2.06</u> | <u>48.6 ± 3.07</u> | 28.7 ± 3.05 | **65.8 ± 2.55** |
| | PhenoCell<sub>all</sub> | <u>18.0 ± 5.00</u> | 17.5 ± 5.26 | 9.7 ± 2.67 | 12.9 ± 1.86 | 16.6 ± 3.37 | 17.7 ± 4.51 | **20.7 ± 3.70** |
| | PhenoCell<sub>imm</sub> | 16.3 ± 6.10 | 16.0 ± 5.07 | 7.9 ± 2.88 | 8.8 ± 1.23 | <u>17.6 ± 3.78</u> | 17.2 ± 4.40 | **18.6 ± 3.84** |

### 3.4 Ablation study

#### Branch-conditional calibration reduces class-wise coverage gaps

To motivate the design choices in our COF module with respect to class-wise coverage, we compare against standard-split CP and graph-based CP from Corbetta et al. (2026) with the same CellBridge representations and linear probing. We measure CovGap, the mean absolute deviation of per-class empirical coverage from the nominal 90% target, in percentage points

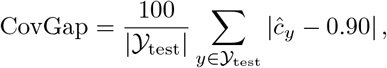

where 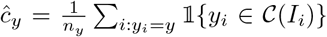 is the empirical coverage for class *y, Y*_test_ contains the classes present in the test set, *n*_y_ denotes their respective test counts, and *C*(*I*_*i*_) is the predicted label set for sample *I*_*i*_. Table 4 establishes that incorporating the ontology reduces CovGap across all tasks, with branch-conditional calibration providing further consistent improvements, especially for undercovered rare classes.

**Table 4:** Effect of conformal prediction strategies on class-wise coverage gaps. CovGap in percentage points (↓) for standard, graph-based, and branch-conditional CP using CellBridge. Values report the mean and standard error, with the latter derived from the standard deviation across the ID sections and estimated via 1000 grouped-bootstrap replicates for OOD datasets. Bold and underline indicate the best and second-best per row, respectively.

| Dataset |  | Standard | Graph | Ours |
| --- | --- | --- | --- | --- |
| ID | HEST | 21.2 ± 2.40 | <u>13.4 ± 1.48</u> | **8.6 ± 0.89** |
| OOD | NuCLS | 18.2 ± 2.43 | <u>10.1 ± 1.72</u> | **4.8 ± 0.65** |
| | PanNuke | 3.6 ± 0.32 | <u>3.3 ± 0.31</u> | **1.5 ± 0.19** |
| | Lizard | 24.1 ± 1.51 | <u>16.8 ± 0.99</u> | **4.6 ± 0.36** |
| | PhenoCell<sub>all</sub> | 18.5 ± 2.19 | <u>7.7 ± 1.44</u> | **5.4 ± 1.11** |
| | PhenoCell<sub>imm</sub> | 22.3 ± 2.80 | <u>7.9 ± 2.26</u> | **4.1 ± 1.09** |

#### scFM choice determines the quality of transferred representations

Having established the benefit of molecular guidance in Table 2, we now examine whether the specific teacher choice matters. Accordingly, we replace TERRA embeddings with those from NOVAE (Blampey et al., 2025) while retaining the same adaptation setup. As shown in Table 5, TERRA yields higher linear-probe macro F1 on every task. The largest gains occur on Lizard, demonstrating that scFM choice substantially affects transfer beyond the pretraining cohort. Crucially, even with NOVAE as the teacher, our model still surpasses the strongest image-only foundation models in Table 2 on datasets with the most complex label spaces, namely NuCLS and PhenoCell_all_, demonstrating the benefits of molecular guidance. These results support TERRA as the teacher model for CellBridge and show that the benefits of multi-modal alignment depend on the representations used to guide adaptation.

**Table 5:**
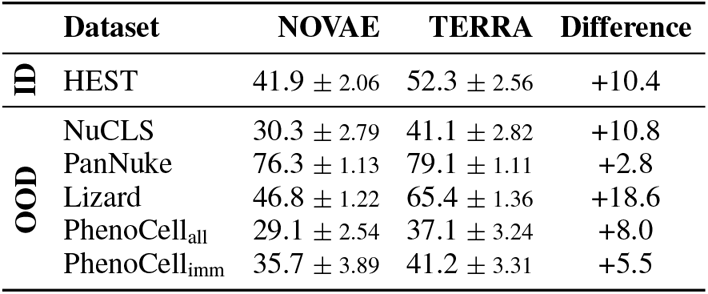
Effect of the single-cell foundation model on transfer. Linear-probe macro F1 (%) for CellBridge pretrained with NOVAE or TERRA. Values report the mean and standard error, with the latter derived from the standard deviation across the ID sections and estimated via 1000 grouped-bootstrap replicates for OOD datasets. Differences are in percentage points.

### 3.5 Applied case study

#### Deriving biological insights with CellBridge

To explore CellBridge’s practical value, we performed a qualitative case study with expert biologists, who jointly reviewed H&E images, reference annotations, model predictions, and reliability scores. Figure 4 illustrates one such example from a breast cancer tissue where two macrophages (orange) are infiltrating a cluster of tumor cells (blue). Although our model misclassifies these macrophages as tumor cells, its COF module identifies the ambiguity and assigns low reliability scores to these exact cells. This confirms that CellBridge is adapted to the complexity of cellular microenvironments.

**Figure 4.**
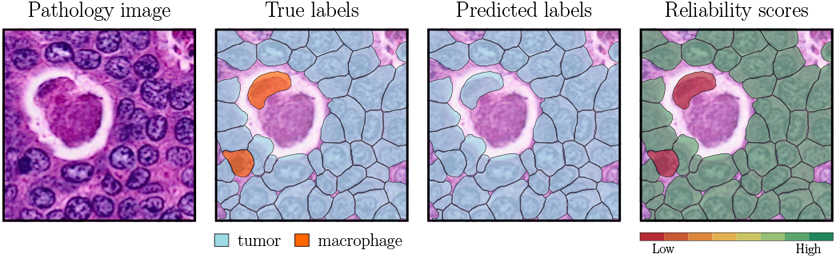
Applied case study. A breast tissue, its ground-truth cell labels, CellBridge predictions, and reliability scores per cell from our COF module. Additional examples in Appendix A.4.

**Figure 5.**
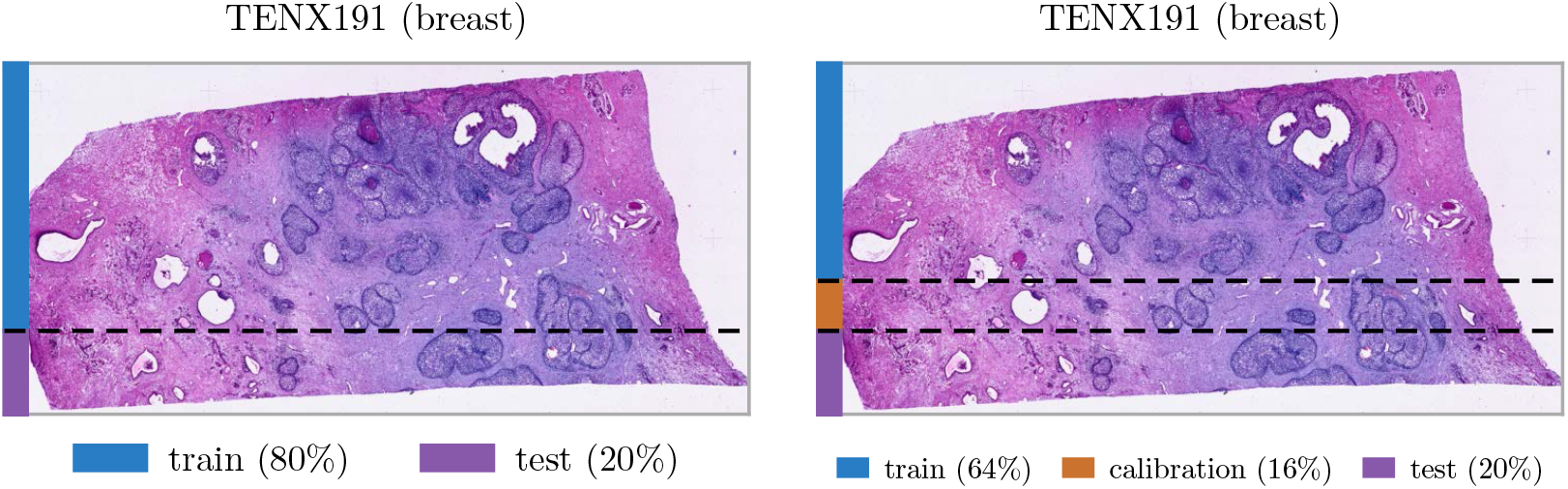
Example spatial splits for downstream analysis (left) and conformal prediction (right).

## 4 Related work

### Pathology foundation models

Large-scale self-supervised pretraining has produced general-purpose foundation models operating on H&E images, such as UNI2-h (Chen et al., 2024) and Virchow2 (Zimmermann et al., 2024). These models learn transferable representations from tissue patches and achieve state-of-the-art performance on region- and slide-level tasks. However, their inputs typically comprise 224 × 224-pixel patches, whose physical field of view depends on image resolution. Such patches often contain multiple cells, and hence, their representations do not explicitly distinguish individual cells, which is required for fine-grained phenotyping.

### Cell-level representation learning

Dedicated cell encoders address the granularity mismatch of patch-level methods by explicitly modeling individual cells. VOLTA (Nakhli et al., 2024) combines two contrastive objectives: a Cell Block aligns augmented views of the same cell while separating different cells and an Environment Block aligns each target cell with its surrounding tissue context. LEMON (Chadoutaud et al., 2026) learns cellular representations through MoCo v3 pretraining on cell-centered H&E images. Although these methods capture cell-level morphology, their image-only pretraining cannot differentiate between molecularly defined phenotypes with similar appearances.

### Molecularly guided representations

The growing availability of paired H&E images and ST data, especially the public HEST-1k (Jaume et al., 2024) dataset, has made it possible to learn joint image–expression spaces. However, the majority of these multi-modal approaches, such as SEAL (Hemker et al., 2026), STORM (Xiang et al., 2026), and STPath (Huang et al., 2025), rely on spot-level ST platforms. In other words, the output expression profiles are aggregated across multiple cells into spots, and therefore, they inherently obscure cellular details. Despite recent exceptions like PAST (Yang et al., 2025), which performs alignment directly at single-cell resolution, and SpatialWhisperer (Schaefer et al., 2026), which relaxes the pairing requirement altogether by using gene expression to bridge H&E images and textual descriptions in a trimodal setting, a critical gap remains. In particular, none of these approaches quantify the reliability of their predictions.

## 5 Conclusion

We presented CellBridge, a framework for fine-grained and reliable cell phenotyping from H&E images. CellBridge leverages a large corpus of paired single-cell observations to align representations from pretrained pathology and ST foundation models, transferring molecular information into the image encoder while requiring only H&E at inference. To account for phenotypic distinctions that cannot be reliably resolved from morphology, our COF module adjusts the granularity of output predictions. Across held-out tissue sections and external datasets, CellBridge substantially improves classification performance, while our COF module flags ambiguous cells. However, several limitations remain. First, CellBridge requires precomputed cell segmentations, although established cell annotators can mitigate this requirement. Second, its performance depends on the quality of the underlying foundation model embeddings, and may therefore benefit from continued advances in both modalities. Third, pretraining is currently constrained by the size and diversity of the HEST-1k Xenium cohort, highlighting the value of larger and more diverse paired datasets. Overall, CellBridge advances the analysis of tissue microenvironments from routine pathology images, while explicitly representing the limits of what can be reliably inferred from morphology alone.

### Ause statement

In this work, we used generative AI tools to:

1. Design or provide feedback on research methodology or experiments. The core ideas are all human-generated.
2. Implement methods, especially to optimize parts of our code for efficiency. All code was reviewed by the authors, who take full responsibility for its correctness.
3. Support qualitative and thematic data analysis, especially in making plots and visualizations.

We did not use generative AI tools to:

1. Clean and reformat datasets.
2. Interpret results.

These required disclosure tasks do not apply to this work:

1. Generate synthetic data sets.
2. Help develop theoretical models or conceptual frameworks.
3. Formulate mathematical claims.
4. Provide critical ingredients for proving mathematical claims.
5. Assist in the writing of proofs.
6. Propose or refine hypotheses.
7. Assist with translation.

Additionally, we used generative AI tools to:

1. Edit manuscript text to improve readability, for example, to find and correct grammar mistakes.
2. Identify relevant literature. All retrieved work was manually reviewed by authors.

We take full responsibility for the final manuscript, including all content produced with the assistance of generative AI.

### Ethics statement

This work uses publicly available histopathology and spatial transcriptomics datasets, with patient data anonymized by the original data providers. Therefore, the authors do not foresee ethical concerns arising from this work.

### Reproducibility statement

Our implementation uses Python 3.12 and PyTorch. We provide the complete training and evaluation pipeline at https://github.com/Lotfollahi-lab/cellbridge, along with a full list of dependencies. All experiments were conducted on a single NVIDIA H200 GPU with 141GB of memory, using a fixed random seed across all stochastic components. The appendix provides the implementation details needed to reproduce our results.

## A Appendix

### A.1 Dataset details

#### HEST-1k

Jaume et al. (2024) introduced this collection of spatial transcriptomic profiles, each aligned to an H&E-stained whole-slide image, and accompanied by curated metadata. Although it spans several assay families, we focus our analysis on human 10x Xenium In Situ sections from v1.3.0, which are imaging-based and provide transcripts assigned to individually segmented cells. This yields 79 sections^2^ spanning 14 organs, with a total of 23.8M cells, and Table 6 reports their organ distribution. We partition these sections into a training set (59 samples and 16M cells) for multi-modal alignment, a validation set (6 samples and 2.8M cells) for model selection and early stopping, and a held-out annotated test set (14 samples and 4M cells) to evaluate cell phenotyping performance.

**Table 6:**
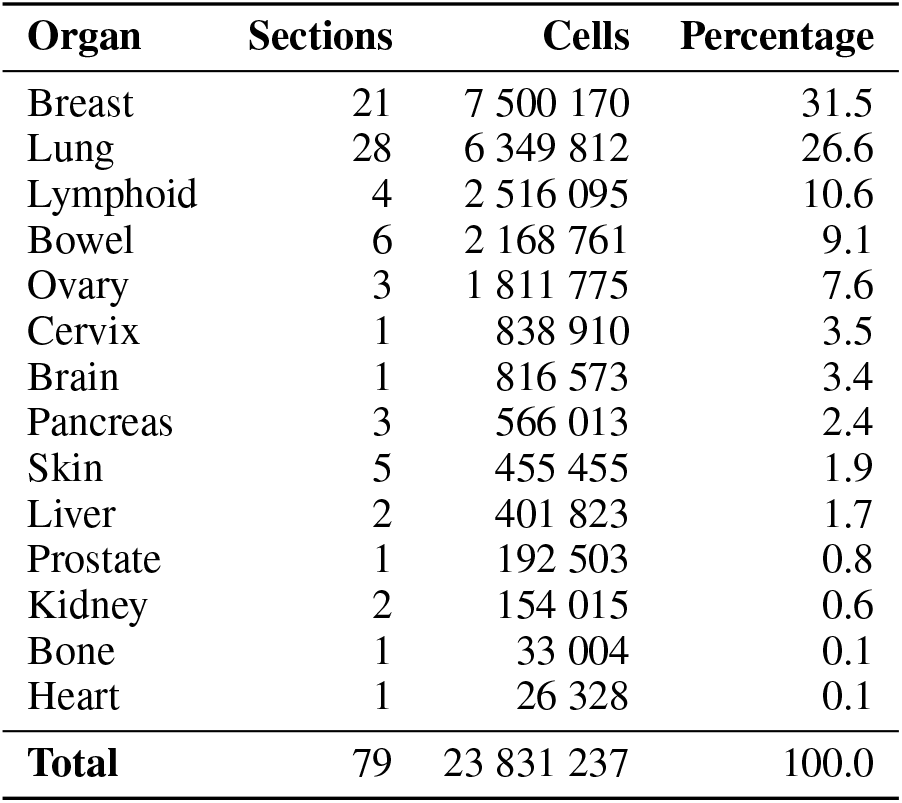
Composition of used HEST sections by organ, ordered by cell count.

| Organ | Sections | Cells | Percentage |
| --- | --- | --- | --- |
| Breast | 21 | 7 500 170 | 31.5 |
| Lung | 28 | 6 349 812 | 26.6 |
| Lymphoid | 4 | 2 516 095 | 10.6 |
| Bowel | 6 | 2 168 761 | 9.1 |
| Ovary | 3 | 1 811 775 | 7.6 |
| Cervix | 1 | 838 910 | 3.5 |
| Brain | 1 | 816 573 | 3.4 |
| Pancreas | 3 | 566 013 | 2.4 |
| Skin | 5 | 455 455 | 1.9 |
| Liver | 2 | 401 823 | 1.7 |
| Prostate | 1 | 192 503 | 0.8 |
| Kidney | 2 | 154 015 | 0.6 |
| Bone | 1 | 33 004 | 0.1 |
| Heart | 1 | 26 328 | 0.1 |
| **Total** | 79 | 23 831 237 | 100.0 |

**Table 7:** Class balance of the source NuCLS and PhenoCell datasets, each ordered by cell count.

| (a) NuCLS. |  |  | (b) PhenoCell. |  |  |
| --- | --- | --- | --- | --- | --- |
| Class | Cells | Percentage | Class | Cells | Percentage |
| Tumor | 21 067 | 35.42 | Tumor cells | 49 045 | 18.46 |
| Lymphocyte | 13 630 | 22.91 | Macrophages / monocytes | 46 186 | 17.39 |
| Fibroblast | 8 620 | 14.49 | T cells | 43 547 | 16.39 |
| Unlabeled | 7 515 | 12.63 | Smooth muscle | 28 926 | 10.89 |
| Plasma cell | 5 562 | 9.35 | Granulocytes | 22 247 | 8.38 |
| Macrophage | 1 354 | 2.28 | Stroma | 21 091 | 7.94 |
| Vascular endothelium | 512 | 0.86 | B cells | 15 176 | 5.71 |
| Ductal epithelium | 498 | 0.84 | Other cells | 13 925 | 5.24 |
| Apoptotic body | 396 | 0.67 | Vasculature / lymphatics | 13 205 | 4.97 |
| Mitotic figure | 229 | 0.38 | Plasma cells | 8 937 | 3.36 |
| Myoepithelium | 54 | 0.09 | Adipocytes | 1 850 | 0.70 |
| Neutrophil | 45 | 0.08 | Nerves | 683 | 0.26 |
| Eosinophil | 3 | 0.01 | Dendritic cells | 438 | 0.16 |
|  |  |  | NK cells | 374 | 0.14 |
| **Total** | **59 485** | **100.0** | **Total** | **265 630** | **100.0** |

### NuCLS

This dataset from Amgad et al. (2022) contains annotated nuclei from breast cancer images, with labels organized at three granularities: a raw set of 13 classes, a main set of 8 classes, and a super set of 5 classes obtained by aggregating the raw labels into broader groups. We use the pathologist-corrected single-rater subset, comprising 59k annotations. Table 7a reports the source label distribution. Since we target fine-grained phenotyping, we adopt the raw labels after discarding unlabeled annotations and partition the dataset into 29k train, 16k validation, and 6k test cells.

The following datasets come from PathoCellBench (Lüscher et al., 2025), a comprehensive bench-mark for evaluating the cell phenotyping capabilities of pathology foundation models.

### PhenoCell

Lüscher et al. (2025) built this dataset from CODEX-imaged FFPE colon carcinoma sections collected at the University Hospital Bern, transferring marker-based cell phenotypes onto the paired H&E fields of view. It provides a 35-patient cohort with 266k annotated cells assigned to 14 classes. We follow the benchmark’s base split, which keeps all fields from the same tissue region in the same partition: 74 fields (185k cells) for training, 16 fields (38k cells) for validation, and 19 fields (43k cells) for testing. We report performance over all 14 classes and over the immune subset of 7 classes. Table 7b similarly reports the resulting label distribution.

### Lizard

Graham et al. (2021) assembled this colon H&E dataset by pooling nuclei annotations from several cohorts. It provides 238 image regions with 469k nuclei labeled as one of six classes. We follow the benchmark’s default three-fold split, which holds out whole regions: 98 regions (139k cells) for training, 70 regions (167k cells) for validation, and 70 regions (162k cells) for testing. Table 8a reports the resulting label distribution.

### PanNuke

Gamper et al. (2020) built this dataset from 256 × 256 H&E patches spanning 19 tissue types, with semi-automatically generated nuclei masks refined by expert review. The release we use contains 7.5k patches and 225k nuclei assigned to five classes. We follow the official fold assignment provided by the benchmark, using fold 3 for training (79k cells), fold 1 for validation (75k cells), and fold 2 for testing (71k cells). Table 8b reports the resulting label distribution.

## A.2 Baseline details

### A.2.1 Patch-level foundation models

#### UNI2-h

The UNI2-h encoder (Chen et al., 2024) is a patch-level FM that scales the original UNI architecture to a ViT-H/14 backbone of 681M parameters. Pretraining follows the DINOv2 objective and utilizes 200M tiles taken from more than 350k H&E- and IHC-stained slides at Mass General Brigham. We use the Trident (Zhang et al., 2025) toolkit for checkpoint loading and extract the class token, yielding a 1536-dimensional representation.

#### Virchow2

The patch-level FM of Virchow2 (Zimmermann et al., 2024) keeps the ViT-H/14 backbone (632M parameters) of its predecessor but adapts the DINOv2 objective for histopathology. Pretraining data comprises 3.1M H&E- and IHC-stained slides from Memorial Sloan Kettering CancerCenter, tiled across four magnifications (5×, 10×, 20×, 40×). Similarly, we use the Trident toolkit and follow the recommended representation by concatenating the class token with the mean of the patch tokens, producing a 2560-dimensional embedding.

### A2.2 Cell-level foundation models

#### VOLTA

Introduced by Nakhli et al. (2024), VOLTA is a cell-level FM that explicitly incorporates the local tissue environment around each target cell with a contrastive objective. It is specifically designed for annotation-free cell phenotyping and was evaluated across more than 800k cells spanning six cancer types. We use the released ovarian checkpoint, from which we extract the raw 512-dimensional representation for each cell.

#### LEMON

Proposed by Chadoutaud et al. (2026), LEMON learns generalizable nuclear morphology representations through a MoCo v3 objective on 10M cropped nuclei. Its backbone is a ViT-S/8 operating on 40 × 40 images, leading to 384-dimensional cell-level embeddings.

### A.2.3 Multi-modal foundation models

#### PAST

This is a multi-modal single-cell foundation model for histopathology and spatial transcriptomics (Yang et al., 2025). It is pretrained on 19.3M paired single-cell image-transcriptome examples across 15 cancer types. For each cell, PAST encodes a 224 × 224 H&E patch with a

ViT-L/16 image encoder, while the transcriptomic branch represents each cell as a ranked sequence of expressed gene tokens and incorporates local spatial context with a graph attention module. We use the released pretrained encoder to extract 1024-dimensional cell-level representations.

#### SpatialWhisperer

This trimodal contrastive model aligns H&E image patches, gene expression profiles, and natural-language cell annotations in a shared embedding space (Schaefer et al., 2026). It leverages the shared gene-expression modality to transfer supervision transitively from image-spatial-transcriptomics pairs and single-cell RNA sequencing-text annotation pairs. We use only the image branch, taking the 2048-dimensional encoder output as our cell representation.^3^

## A.3 Implementation details

### A.3.1 Preprocessing

For HEST-1k, we reconstruct cell-by-gene count matrices from the Xenium transcript files after excluding unassigned transcripts, control probes, and transcript calls with quality scores below 20. We aggregate the retained transcripts by cell and gene, map gene symbols to TERRA’s Ensembl vocabulary, and remove genes detected in fewer than 10 cells and cells with fewer than 10 detected genes. We then construct a 10-nearest-neighbor graph within each section using native Xenium coordinates for TERRA tokenization. The resulting TERRA embeddings are matched to the corresponding H&E inputs by cell identifier. For each cell, we extract an H&E crop spanning 112 × 112 *µ*m around its center and resize it to 224 × 224 pixels using bilinear interpolation, yielding a resolution of 0.5 *µ*m/pixel. Regions extending beyond slide boundaries are padded with white pixels. These crops serve as inputs to UNI2-h for image embedding extraction.

For external datasets, we locate cells using the released annotations and extract a 224 × 224 source-pixel crop centered on each cell, padding regions outside the image boundaries with white pixels.

### A.3.2 Pretraining

After preprocessing, we train on 16 335 536 paired cells from 59 HEST sections. Validation uses randomly sampled 5 000 cells from each of six separate sections, giving 30 000 cells. We keep the pretrained encoder weights fixed and optimize only the LoRA layers (Hu et al., 2022) and image projection head, containing 294 912 and 593 280 trainable parameters, respectively. Training uses AdamW (Loshchilov & Hutter, 2019), learning rates of 10^−4^ for LoRA and 3 × 10^−4^ for the projection head, and weight decay 10^−4^, with a batch size of 16. We select the checkpoint with the lowest validation loss.

### A.3.3 Evaluation: downstream analysis

#### Linear probing

We keep all image encoders frozen and fit an _2_-regularized logistic regression classifier on standardized embeddings. For HEST, we reserve a spatially disjoint 20% of cells in each section for testing and select the regularization parameter *C* ∈ 0.1, 1, 10} by three-fold stratified cross-validation on the remaining 80% (see Figure 5 left), using macro F1. We then refit the classifier on all training cells. For external benchmarks, we select *C* using the provided validation split and refit on the combined training and validation sets before testing.

#### kNN

We also evaluate frozen embeddings using a *k*-nearest-neighbor classifier with *k* = 20, cosine distance, and uniform voting. For HEST, the reference set contains the 80% training partition, whereas for OOD datasets, it combines the training and validation partitions. We use the same test sets as for linear probing and keep *k* fixed across datasets and representations.

### A.3.4 Evaluation: conformal prediction

For conformal prediction, we keep classifier fitting and calibration separate. HEST sections are partitioned spatially into approximately 64% training, 16% calibration, and 20% testing cells, preserving the same test stripe (see Figure 5 right). For OOD datasets, the provided training, validation, and test splits serve as training, calibration, and testing sets, respectively. In both cases, we adapt the linear probing setting described above, select *C* by three-fold stratified cross-validation within the training set, and fit the classifier without using calibration cells. We then calibrate prediction sets at *α* = 0.1, corresponding to a nominal coverage target of 90%. Note that we map labels with the pinned Cell Ontology snapshot dated June 8^th^ 2026 and exclude unmapped phenotypes as well as those represented by fewer than 20 cells. The supplementary material provides dataset-specific taxonomy mappings and excluded labels in YAML files, ontology visualizations in PDF format, and the implementation of the branching rule.

## A.4 Additional results

This section reports additional metrics that complement the main results. We provide balanced accuracy and macro AUROC for both cross-section generalization (Figure 6) and OOD generalization (Tables 10 and 11). The corresponding empirical coverage results of Table 3 are given in Table 12. We observe that all methods satisfy the 90% coverage target under our COF module. Across all these metrics, the results are consistent with our previous findings. Figures 7, 8, 9, 10, 11, 12, and 13 provide further qualitative visualizations with expert explanations as captions.

**Table 8:** Class balance of the source Lizard and PanNuke datasets, each ordered by cell count.

| (a) Lizard. |  |  | (b) PanNuke. |  |  |
| --- | --- | --- | --- | --- | --- |
| Class | Cells | Percentage | Class | Cells | Percentage |
| Epithelial | 231 526 | 49.38 | Neoplastic | 93 675 | 41.60 |
| Connective tissue | 105 471 | 22.49 | Connective / soft tissue | 59 798 | 26.56 |
| Lymphocyte | 97 666 | 20.83 | Inflammatory | 36 913 | 16.39 |
| Plasma | 26 499 | 5.65 | Epithelial | 31 614 | 14.04 |
| Neutrophil | 4 465 | 0.95 | Dead | 3 171 | 1.41 |
| Eosinophil | 3 246 | 0.69 |  |  |  |
| **Total** | **468 873** | **100.0** | **Total** | **225 171** | **100.0** |

**Table 9:**
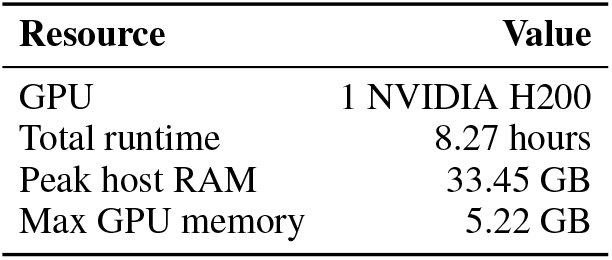
Time and memory requirements for pretraining.

**Table 10:** Balanced accuracy (%) scores comparing CellBridge against baseline FMs using both linear and kNN probes. Values denote the mean and standard error computed across 1000 grouped-bootstrap replicates of the fixed test sets. SW stands for SpatialWhisperer. Bold and underline indicate the best and second-best per row, respectively.

|  |  | Patch-level FMs |  | Cell-level FMs |  | Multi-modal FMs |  | Ours |
| --- | --- | --- | --- | --- | --- | --- | --- | --- |
|  |  | UNI2-h | Virchow2 | VOLTA | LEMON | PAST | SW | CellBridge |
| Linear Probe | NuCLS | 26.4 ± 1.52 | 27.2 ± 2.64 | 25.5 ± 2.26 | 25.2 ± 2.48 | 26.8 ± 2.04 | 27.0 ± 2.54 | **40.7 ± 2.60** |
| | PanNuke | 71.8 ± 1.52 | 71.3 ± 1.70 | 54.7 ± 0.85 | 66.5 ± 1.17 | <u>73.2 ± 1.28</u> | 73.0 ± 1.29 | **78.3 ± 1.24** |
| | Lizard | 34.3 ± 1.11 | 33.9 ± 1.56 | 54.9 ± 1.17 | **65.3 ± 1.07** | 45.3 ± 1.07 | 36.2 ± 0.93 | <u>62.1 ± 1.31</u> |
| | PhenoCell<sub>all</sub> | <u>25.2 ± 1.98</u> | 24.7 ± 2.05 | 16.1 ± 0.89 | 22.1 ± 1.26 | 22.9 ± 1.61 | 24.7 ± 1.93 | **35.9 ± 2.76** |
| | PhenoCell<sub>imm</sub> | <u>35.9 ± 4.03</u> | 33.3 ± 3.96 | 23.4 ± 1.27 | 30.5 ± 1.92 | 28.7 ± 2.04 | 32.8 ± 3.27 | **41.0 ± 3.52** |
| kNN | NuCLS | 22.7 ± 0.93 | 23.1 ± 1.39 | 21.8 ± 0.67 | <u>23.9 ± 1.16</u> | 20.1 ± 1.33 | 23.5 ± 1.09 | **33.4 ± 1.94** |
| | PanNuke | 73.9 ± 1.44 | 74.2 ± 1.38 | 46.0 ± 0.47 | 64.3 ± 1.26 | <u>75.9 ± 1.19</u> | 74.1 ± 1.47 | **80.8 ± 1.07** |
| | Lizard | 28.4 ± 1.11 | 28.0 ± 1.04 | 40.6 ± 0.61 | 46.2 ± 1.09 | 33.6 ± 0.78 | 29.3 ± 1.10 | **58.2 ± 1.36** |
| | PhenoCell<sub>all</sub> | <u>22.6 ± 1.89</u> | 20.0 ± 1.45 | 14.0 ± 0.79 | 17.2 ± 1.09 | 18.3 ± 1.26 | 22.0 ± 1.83 | **34.6 ± 2.59** |
| | PhenoCell<sub>imm</sub> | 30.2 ± 3.43 | 26.9 ± 2.43 | 21.4 ± 1.42 | 24.6 ± 1.66 | 23.5 ± 1.84 | <u>30.4 ± 3.32</u> | **38.9 ± 2.95** |

**Table 11:** Macro AUROC (%) scores comparing CellBridge against baseline FMs using both linear and kNN probes. Values denote the mean and standard error computed across 1000 grouped-bootstrap replicates of the fixed test sets. SW stands for SpatialWhisperer. Bold and underline indicate the best and second-best per row, respectively.

|  |  | Patch-level FMs |  | Cell-level FMs |  | Multi-modal FMs |  | Ours |
| --- | --- | --- | --- | --- | --- | --- | --- | --- |
|  |  | UNI2-h | Virchow2 | VOLTA | LEMON | PAST | SW | CellBridge |
| Linear Probe | NuCLS | 85.6 ± 2.44 | 83.0 ± 2.28 | <u>86.3 ± 2.39</u> | 79.0 ± 3.72 | 73.9 ± 5.53 | 79.2 ± 3.35 | **92.0 ± 1.26** |
| | PanNuke | 94.0 ± 0.22 | 94.2 ± 0.21 | 87.1 ± 0.36 | 92.0 ± 0.22 | <u>94.9 ± 0.17</u> | 94.3 ± 0.23 | **96.6 ± 0.13** |
| | Lizard | 85.5 ± 1.00 | 83.9 ± 1.15 | 93.9 ± 0.31 | <u>96.2 ± 0.26</u> | 91.5 ± 0.64 | 84.6 ± 1.01 | **96.7 ± 0.30** |
| | PhenoCell<sub>all</sub> | <u>76.4 ± 1.59</u> | 75.8 ± 1.76 | 69.2 ± 1.48 | 74.1 ± 1.39 | 75.1 ± 1.77 | 75.1 ± 1.45 | **82.2 ± 1.80** |
| | PhenoCell<sub>imm</sub> | 73.3 ± 2.87 | 74.1 ± 3.35 | 69.1 ± 2.33 | <u>74.8 ± 1.90</u> | 72.0 ± 2.23 | 71.9 ± 2.30 | **80.5 ± 2.68** |
| kNN | NuCLS | 77.4 ± 1.67 | 72.3 ± 2.20 | 75.8 ± 2.27 | <u>78.1 ± 2.51</u> | 68.7 ± 2.80 | 76.2 ± 1.48 | **86.8 ± 1.49** |
| | PanNuke | 93.2 ± 0.28 | 93.2 ± 0.31 | 80.4 ± 0.40 | 90.1 ± 0.45 | <u>94.8 ± 0.24</u> | 93.4 ± 0.27 | **96.3 ± 0.17** |
| | Lizard | 71.1 ± 1.20 | 70.6 ± 1.16 | 84.2 ± 0.53 | <u>89.5 ± 0.62</u> | 77.5 ± 1.26 | 71.9 ± 1.20 | **92.5 ± 0.51** |
| | PhenoCell<sub>all</sub> | <u>69.4 ± 1.67</u> | 68.2 ± 1.51 | 60.7 ± 1.26 | 64.1 ± 1.21 | 67.1 ± 1.73 | 69.2 ± 1.62 | **75.5 ± 1.61** |
| | PhenoCell<sub>imm</sub> | <u>70.4 ± 3.04</u> | 67.9 ± 2.81 | 62.5 ± 2.43 | 66.1 ± 2.06 | 66.6 ± 2.95 | 70.2 ± 2.89 | **76.4 ± 2.27** |

**Table 12:**
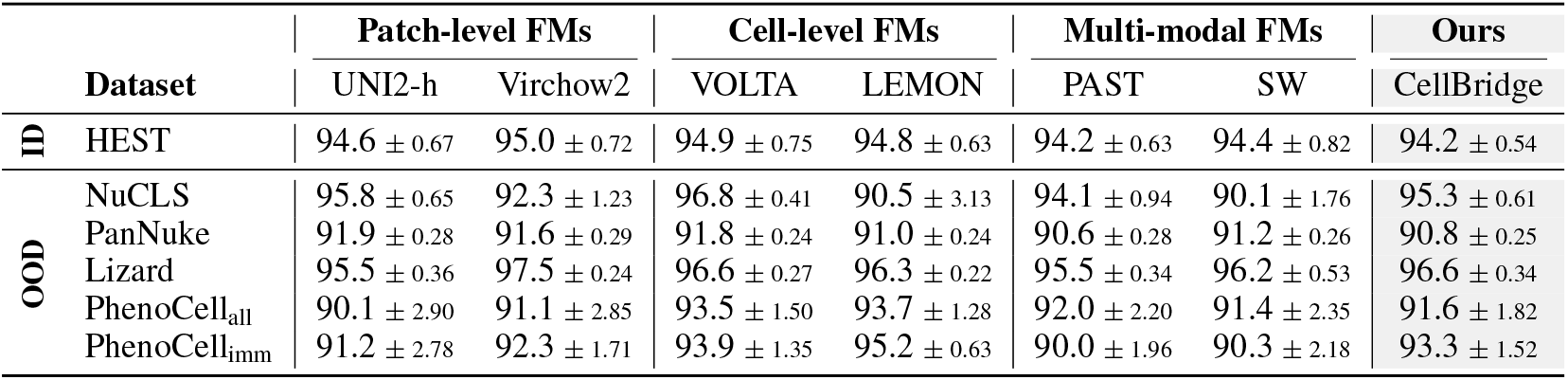
Marginal coverage (%) with target of 90%. Complements Table 3 of the main paper.

**Figure 6.**
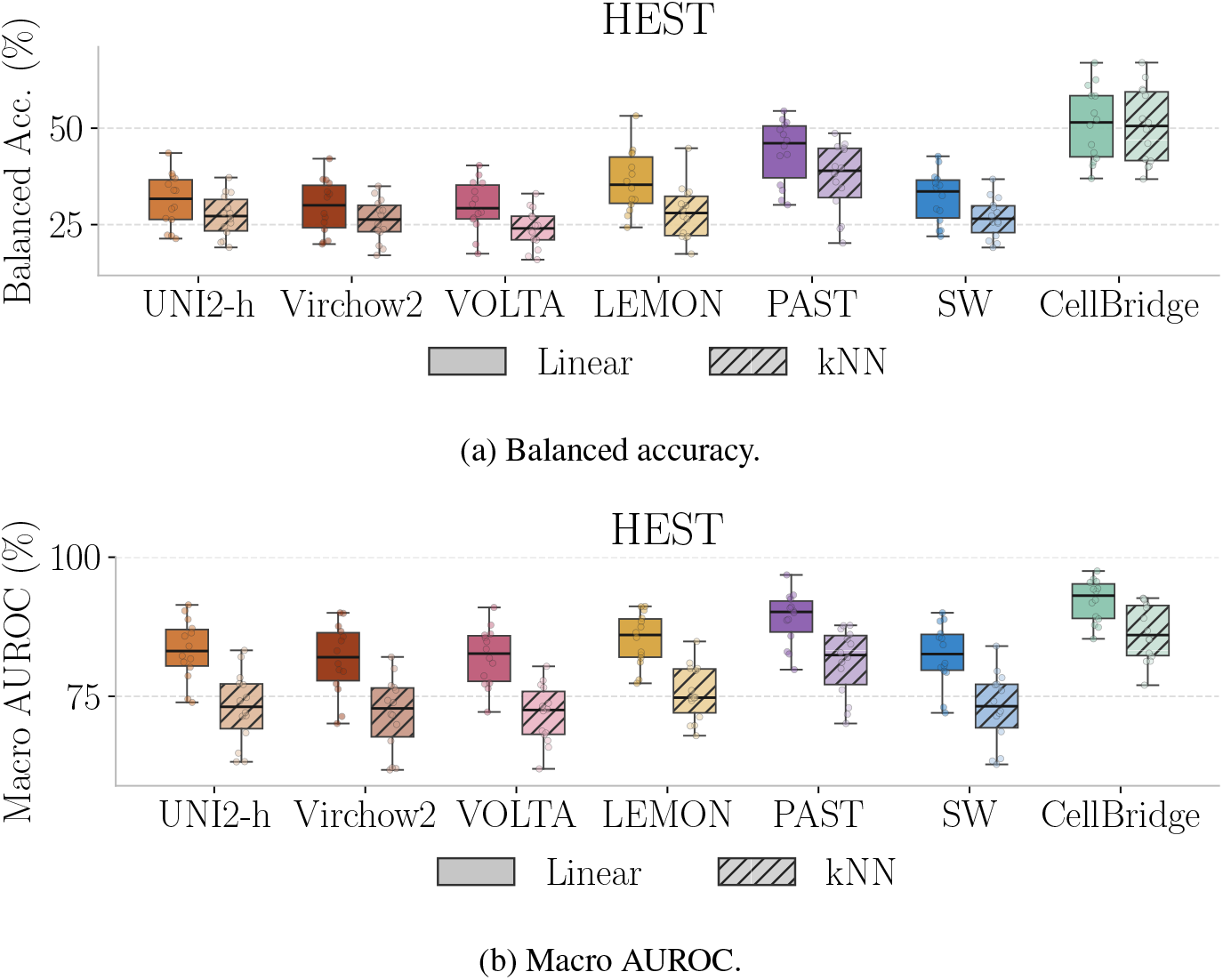
Analogues of Figure 2 for (a) balanced accuracy and (b) macro AUROC.

**Figure 7.**
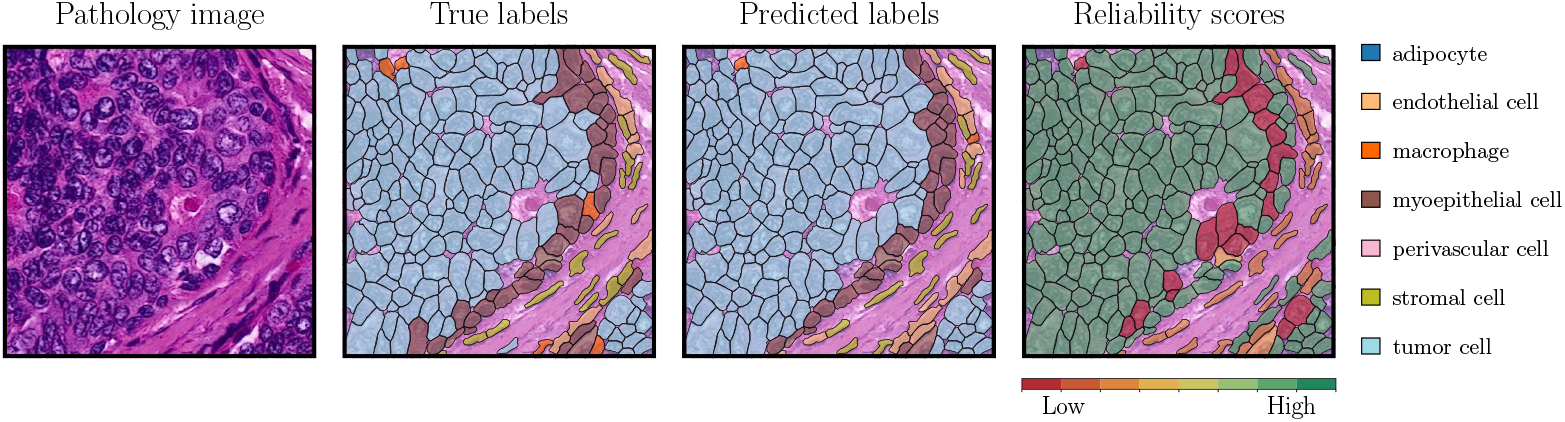
A breast cancer region interpreted by the biologists as a milk duct containing tumor cells, surrounded by an intact myoepithelial layer. Notably, the experts observed that several reference-annotated myoepithelial cells morphologically resembled tumor cells, suggesting the cancer may have originated from the myoepithelium. CellBridge successfully recovers this general spatial organization from images alone, while critically assigning lower reliability scores to these ambiguous cells near the tumor–stromal boundary.

**Figure 8.**
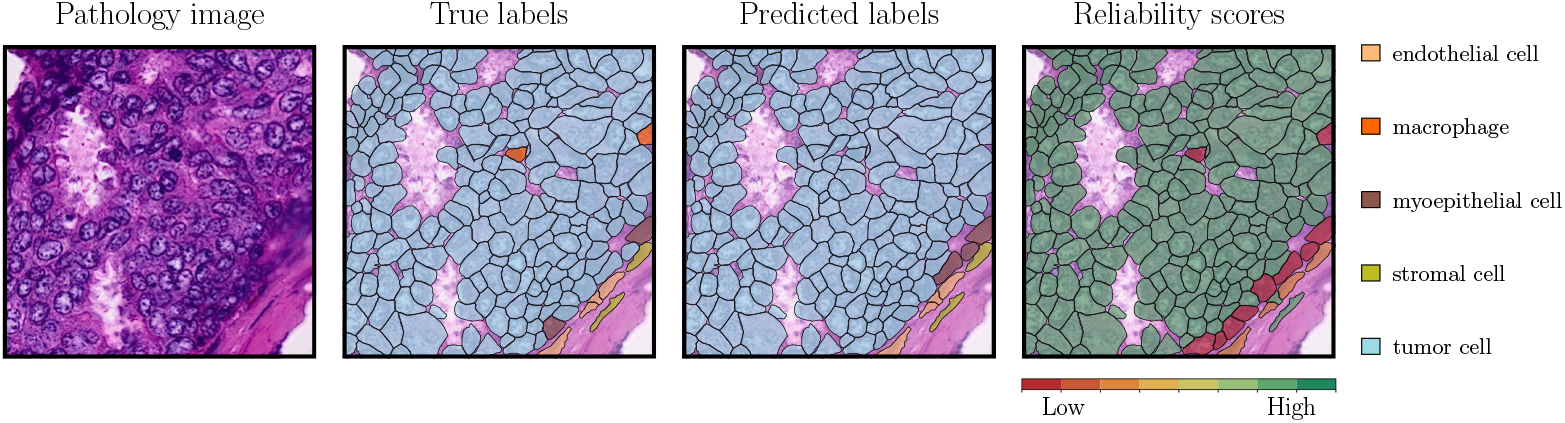
The dominant class is visually well separated: tumor cells have enlarged irregular nuclei and form cohesive nests, while the stromal band at the lower right consists of elongated nuclei aligned with the collagen. The model recovers this boundary and predicts at the leaf level across both regions. The five errors are isolated non-tumor cells surrounded by tumor, where local context is uninformative. All receive near-zero reliability, so they are flagged rather than asserted.

**Figure 9.**
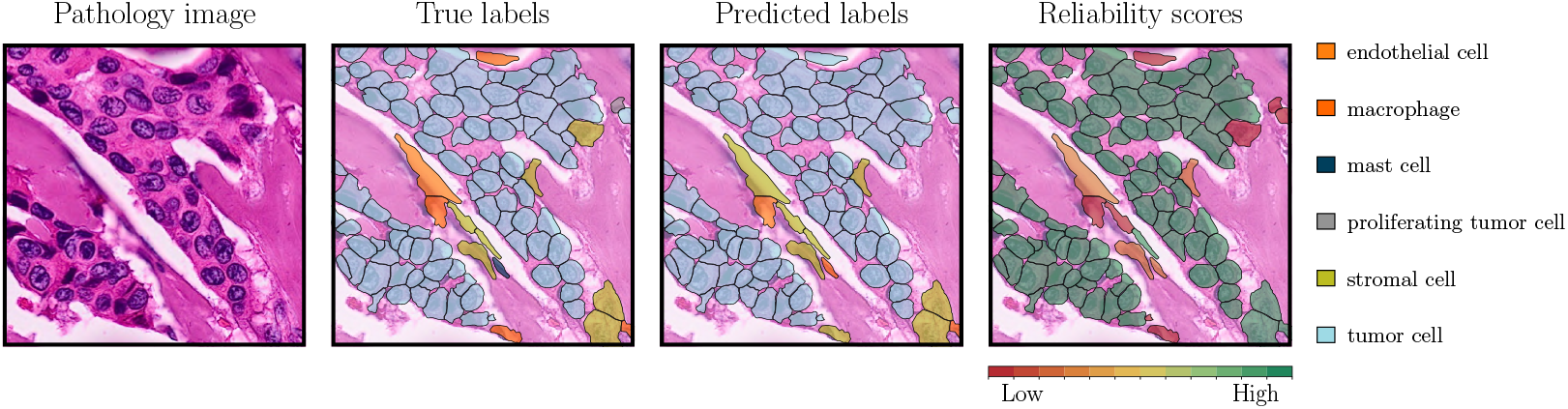
Tumor cells form thin strands between thick collagen bundles, and six classes are present. CellBridge recovers the strand geometry, predicting tumor cells within the strands at the leaf level with high reliability. Errors concentrate on the elongated nuclei compressed between collagen fibers, where endothelial, stromal, and mast cells share a similar spindle-shaped profile. Several of these are predicted as stromal cells, and the COF module assigns them low reliability rather than committing to a specific label.

**Figure 10.**
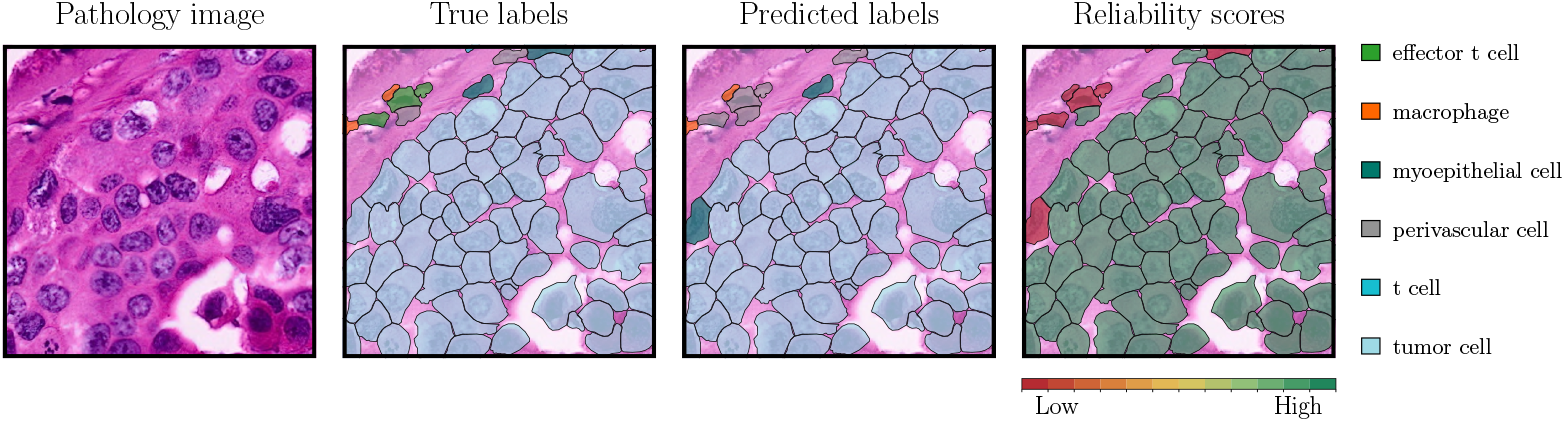
A sheet of tumor cells meets a thin band of immune and perivascular cells along the upper edge, and the model recovers that boundary. Its errors are confined to that band, where T cells, effector T cells, and macrophages are small round cells distinguished in practice by marker expression rather than appearance, and the model assigns them near-zero reliability rather than committing.

**Figure 11.**
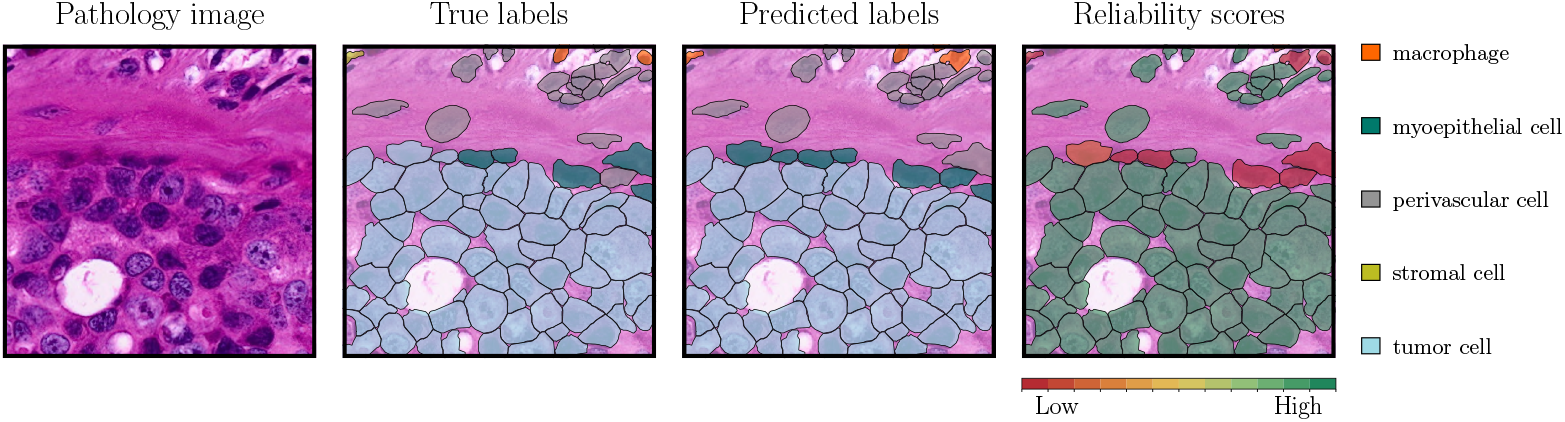
Myoepithelial cells lie along the interface between the tumor-rich region and surrounding connective tissue, consistent with their position around epithelial structures. Confusion between tumor and myoepithelial cells therefore blurs a biologically meaningful boundary. CellBridge assigns low reliability to several cells along this interface, indicating a requirement for closer review.

**Figure 12.**
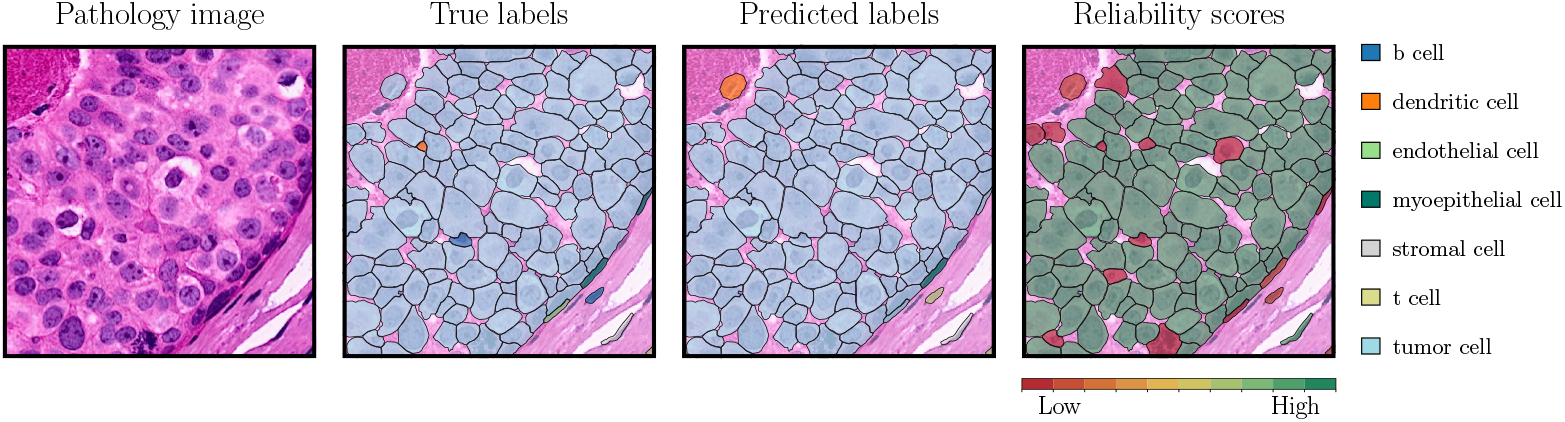
CellBridge recovers the dominant tumor population but misses scattered immune cells, including a dendritic cell and a B cell. Conversely, a tumor cell near the upper-left corner is predicted as dendritic. However, these errors are assigned low reliability scores, showing the use case value.

**Figure 13.**
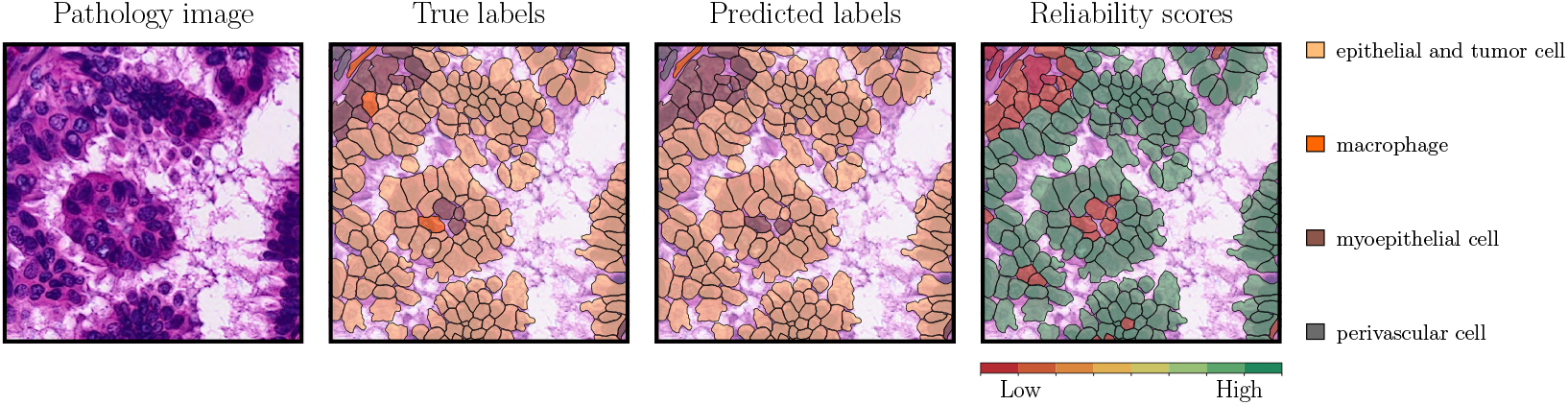
Macrophages lie within and beside groups of epithelial and tumor cells, demonstrating that immune cells are present in regions dominated by epithelial structures. CellBridge captures the major cell groups but misclassifies some macrophages, potentially obscuring their distribution within the tissue. Low reliability highlights these mixed neighborhoods.

## Footnotes

1 Cell segmentation is outside the scope of this work.

2 TENX143 is excluded due to memory constraints

3 The released checkpoint exposes 2048-dimensional features at this layer, as opposed to the reported 512-dimensions in the respective paper.

## References

Mohamed Amgad, Lamees A Atteya, Hagar Hussein, Kareem Hosny Mohammed, Ehab Hafiz, Maha A T Elsebaie, Ahmed M Alhusseiny, Mohamed Atef AlMoslemany, Abdelmagid M Elmatboly, Philip A Pappalardo, Rokia Adel Sakr, Pooya Mobadersany, Ahmad Rachid, Anas M Saad, Ahmad M Alkashash, Inas A Ruhban, Anas Alrefai, Nada M Elgazar, Ali Abdulkarim, Abo-Alela Farag, Amira Etman, Ahmed G Elsaeed, Yahya Alagha, Yomna A Amer, Ahmed M Raslan, Menatalla K Nadim, Mai A T Elsebaie, Ahmed Ayad, Liza E Hanna, Ahmed Gadallah, Mohamed Elkady, Bradley Drumheller, David Jaye, David Manthey, David A Gutman, Habiba Elfandy, and Lee A D Cooper. NuCLS: A scalable crowdsourcing approach and dataset for nucleus classification and segmentation in breast cancer. GigaScience, 11, 2022. ISSN 2047-217X.

Sebastian Birk, Mohammad Vali Sanian, Amirhossein Vahidi, Samuel Ogden, Daniyal J Jafree, Adib Miraki Feriz, Carlo Leonardi, Arpit Merchant, Zichen He, Lloyd Steele, et al. Multi-scale modeling of human tissues from spatial transcriptomics with TERRA. bioRxiv, 2026.

Quentin Blampey, Hakim Benkirane, Nadège Bercovici, Kevin Mulder, Grégoire Gessain, Florent Ginhoux, Fabrice André, and Paul-Henry Cournède. Novae: a graph-based foundation model for spatial transcriptomics data. Nature Methods, 22(12):2539–2550, 2025.

Gabriele Campanella, Shengjia Chen, Manbir Singh, Ruchika Verma, Silke Muehlstedt, Jennifer Zeng, Aryeh Stock, Matt Croken, Brandon Veremis, Abdulkadir Elmas, et al. A clinical benchmark of public self-supervised pathology foundation models. Nature Communications, 16(1): 3640, 2025.

Loïc Chadoutaud, Alice Blondel, Hana Feki, Jacqueline Fontugne, Emmanuel Barillot, and Thomas Walter. LEMON: a foundation model for nuclear morphology in computational pathology. arXiv preprint arXiv:2603.25802, 2026.

Richard J Chen, Tong Ding, Ming Y Lu, Drew F.K. Williamson, Guillaume Jaume, Andrew H Song, Bowen Chen, Andrew Zhang, Daniel Shao, Muhammad Shaban, et al. Towards a general-purpose foundation model for computational pathology. Nature Medicine, 30(3):850–862, 2024.

Daniela Corbetta, Livio Finos, Ludwig Geistlinger, and Davide Risso. Conformal inference for cell-type annotation with graph-structured constraints. Journal of the Royal Statistical Society Series C: Applied Statistics, pp. qlag037, 2026.

Tiffany Ding, Anastasios Angelopoulos, Stephen Bates, Michael Jordan, and Ryan J Tibshirani. Class-conditional conformal prediction with many classes. In NeurIPS, 2023.

Jevgenij Gamper, Navid Alemi Koohbanani, Ksenija Benes, Simon Graham, Mostafa Jahanifar, Syed Ali Khurram, Ayesha Azam, Katherine Hewitt, and Nasir Rajpoot. PanNuke dataset extension, insights and baselines. arXiv preprint arXiv:2003.10778, 2020.

Simon Graham, Mostafa Jahanifar, Ayesha Azam, Mohammed Nimir, Yee-Wah Tsang, Katherine Dodd, Emily Hero, Harvir Sahota, Atisha Tank, Ksenija Benes, Noorul Wahab, Fayyaz Minhas, Shan E Ahmed Raza, Hesham El Daly, Kishore Gopalakrishnan, David Snead, and Nasir Rajpoot. Lizard: A large-scale dataset for colonic nuclear instance segmentation and classification. In ICCVW, pp. 684–693, 2021.

Konstantin Hemker, Andrew H. Song, Cristina Almagro-Pérez, Guillaume Jaume, Sophia J. Wagner, Anurag Vaidya, Nikola Simidjievski, Mateja Jamnik, and Faisal Mahmood. Towards spatial transcriptomics-driven pathology foundation models. arXiv preprint arXiv:2602.14177, 2026.

Arbel Hizmi, Artemii Bakulin, Shai Bagon, and Nir Yosef. Cross-modal knowledge distillation from spatial transcriptomics to histology. arXiv preprint arXiv:2604.09076, 2026.

Edward J Hu, Yelong Shen, Phillip Wallis, Zeyuan Allen-Zhu, Yuanzhi Li, Shean Wang, Lu Wang, and Weizhu Chen. LoRA: Low-rank adaptation of large language models. In ICLR, 2022.

Tinglin Huang, Tianyu Liu, Mehrtash Babadi, Rex Ying, and Wengong Jin. STPath: a generative foundation model for integrating spatial transcriptomics and whole-slide images. npj Digital Medicine, 8(1):659, 2025.

Guillaume Jaume, Paul Doucet, Andrew H Song, Ming Y Lu, Cristina Almagro-Pérez, Sophia J Wagner, Anurag J Vaidya, Richard J Chen, Drew FK Williamson, Ahrong Kim, et al. HEST-1k: A dataset for spatial transcriptomics and histology image analysis. In NeurIPS, 2024.

Ilya Loshchilov and Frank Hutter. Decoupled weight decay regularization. In ICLR, 2019.

Jérôme Lüscher, Nora Koreuber, Jannik Franzen, Fabian H Reith, Claudia Winklmayr, Elias Baumann, Christian M Schürch, Dagmar Kainmüller, and Josef Lorenz Rumberger. PathoCellBench: A comprehensive benchmark for cell phenotyping. In MICCAI, 2025.

Ramin Nakhli, Katherine Rich, Allen Zhang, Amirali Darbandsari, Elahe Shenasa, Amir Hadjifaradji, Sidney Thiessen, Katy Milne, Steven J.M. Jones, Jessica N McAlpine, et al. VOLTA: an environment-aware contrastive cell representation learning for histopathology. Nature Communications, 15(1):3942, 2024.

Florent Peyraud, Jean-Philippe Guegan, Lucile Vanhersecke, Maxime Brunet, Diego Teyssonneau, Lola-Jade Palmieri, Alban Bessede, and Antoine Italiano. Tertiary lymphoid structures and cancer immunotherapy: From bench to bedside. Med, 6(1):100546, 2025.

Alec Radford, Jong Wook Kim, Chris Hallacy, Aditya Ramesh, Gabriel Goh, Sandhini Agarwal, Girish Sastry, Amanda Askell, Pamela Mishkin, Jack Clark, et al. Learning transferable visual models from natural language supervision. In ICML, 2021.

Moritz Schaefer, Zoe Piran, Nils Philipp Walter, Animesh Awasthi, Christoph Bock, Jure Leskovec, and Zinaida Good. Transitive representation learning enhances histopathology annotation. In ICML, 2026.

Shawn Zheng Kai Tan, Aleix Puig-Barbe, Damien Goutte-Gattat, Caroline Eastwood, Brian Aevermann, Alida Avola, James P Balhoff, Ismail Ugur Bayindir, Jasmine Belfiore, Anita Reane Caron, et al. The Cell Ontology in the age of single-cell omics. Scientific Data, 13:946, 2026.

Gbègninougbo Aurel Davy Tchokponhoue, Sevda Üğüt, Ali Idri, Dorina Thanou, and Pascal Frossard. Uncertainty estimation in pathology foundation models via deep mutual learning. arXiv preprint arXiv:2606.30020, 2026.

Vladimir Vovk, David Lindsay, Ilia Nouretdinov, and Alex Gammerman. Mondrian confidence machine. Technical report, Royal Holloway, University of London, 2003.

Vladimir Vovk, Alexander Gammerman, and Glenn Shafer. Algorithmic learning in a random world. Springer, 2005.

Qingjing Wang, Xueting Shao, Yuxuan Zhang, Miaojin Zhu, Frederick X.C. Wang, Jianjian Mu, Jiaxuan Li, Hangping Yao, and Keda Chen. Role of tumor microenvironment in cancer progression and therapeutic strategy. Cancer Medicine, 12(10):11149–11165, 2023.

Jinxi Xiang, Siyu Hou, Yuchen Li, Ryan Quinton, Xiaoming Zhang, Feyisope Eweje, Xiangde Luo, Yijiang Chen, Zhe Li, Colin Bergstrom, et al. A multimodal foundation model of spatial transcriptomics and histology for biological discovery and clinical prediction. arXiv preprint arXiv:2604.03630, 2026.

Changchun Yang, Haoyang Li, Yushuai Wu, Yilan Zhang, Yifeng Jiao, Yu Zhang, Rihan Huang, Yuan Cheng, Yuan Qi, Xin Guo, et al. PAST: A multimodal single-cell foundation model for histopathology and spatial transcriptomics in cancer. arXiv preprint arXiv:2507.06418, 2025.

Xinyu Yuan, Zhihao Zhan, Zuobai Zhang, Manqi Zhou, Jianan Zhao, Boyu Han, Yue Li, and Jian Tang. Cell-Ontology guided transcriptome foundation model. In NeurIPS, 2024.

Andrew Zhang, Guillaume Jaume, Anurag Vaidya, Tong Ding, and Faisal Mahmood. Accelerating data processing and benchmarking of AI models for pathology. arXiv preprint arXiv:2502.06750, 2025.

Eric Zimmermann, Eugene Vorontsov, Julian Viret, Adam Casson, Michal Zelechowski, George Shaikovski, Neil Tenenholtz, James Hall, David Klimstra, Razik Yousfi, et al. Virchow2: Scaling self-supervised mixed magnification models in pathology. arXiv preprint arXiv:2408.00738, 2024.

